# Modulation of triple artemisinin-based combination therapy pharmacodynamics by *Plasmodium falciparum* genotype

**DOI:** 10.1101/2020.07.03.187039

**Authors:** Megan R. Ansbro, Zina Itkin, Lu Chen, Gergely Zahoranszky-Kohalmi, Chanaki Amaratunga, Olivo Miotto, Tyler Peryea, Charlotte V. Hobbs, Seila Suon, Juliana M. Sá, Arjen M. Dondorp, Rob W. van der Pluijm, Thomas E. Wellems, Anton Simeonov, Richard T. Eastman

## Abstract

The first-line treatments for uncomplicated *Plasmodium falciparum* malaria are artemisinin-based combination therapies (ACTs), consisting of an artemisinin derivative combined with a longer acting partner drug. However, the spread of *P. falciparum* with decreased susceptibility to artemisinin and partner drugs presents a significant challenge to malaria control efforts. To stem the spread of drug resistant parasites, novel chemotherapeutic strategies are being evaluated, including the implementation of triple artemisinin-based combination therapies (TACTs). Currently, there is limited knowledge on the pharmacodynamics and pharmacogenetic interactions of proposed TACT drug combinations. To evaluate these interactions, we established an *in vitro* high-throughput process for measuring the drug dose-response to three distinct antimalarial drugs present in a TACT. Sixteen different TACT combinations were screened against fifteen parasite lines from Cambodia, with a focus on parasites with differential susceptibilities to piperaquine and artemisinins. Analysis revealed drug-drug interactions unique to specific genetic backgrounds, including antagonism between piperaquine and pyronaridine associated with gene amplification of *plasmepsin II/III*, two aspartic proteases that localize to the parasite digestive vacuole. From this initial study, we identified parasite genotypes with decreased susceptibility to specific TACTs, as well as potential TACTs that display antagonism in a genotype-dependent manner. Our assay and analysis platform can be further leveraged to inform drug implementation decisions and evaluate next-generation TACTs.

**One Sentence Summary:** *In vitro* process to evaluate triple-drug combinations for prioritizing antimalarial combinations for *in vivo* evaluation.

## Introduction

Recent efforts driving significant reductions in malaria-related morbidity and mortality have relied on long-lasting insecticide-treated bed nets, indoor residual insecticide spraying, rapid diagnostic tests, and artemisinin-based combination therapies (ACTs) (*1*). ACTs are the first-line antimalarial therapy in almost all endemic countries owing to their high efficacy, safety, tolerability, and reduction of transmissibility. ACTs are comprised of two components: a semi-synthetic artemisinin (ART) derivative and a partner drug. ACTs benefit from the ability of the artemisinin component to rapidly reduce the parasite biomass, leaving few parasites to be cleared by partner drugs that possess longer half-lives, and thereby reduce the parasite pool from which resistance can emerge.

However, the effectiveness of ACTs is threatened by the emergence of parasites with decreased susceptibility to the ART derivatives and resistance to several ACT partner drugs. Mutations in the PfKelch13 propeller domain associate with delayed *in vivo* parasite clearance (*2*) that attenuates the efficacy of ART drugs against the ring-stage of the parasite’s erythrocytic life cycle (*3-5*). However, ACTs remain effective, even in regions with high prevalence of parasites harboring *kelch13* mutations, if the partner drug remains efficacious. Nonetheless, any decrease in ART efficacy leaves a higher parasite burden to be eliminated by the partner compound, increasing the potential for parasites to acquire resistance to the partner drug (*6*). Resistance to the ACT partner drugs is associated with higher incidence of ACT clinical treatment failure (*7*). ACTs approved by the World Health Organization (WHO) include artemether+lumefantrine (AL), artesunate+amodiaquine (ASAQ), artesunate+mefloquine (ASM), artesunate+pyronaridine (ASP), and dihydroartemisinin+piperaquine (DP). Except for pyronaridine (PYR), which has not been widely used clinically, all other partner drugs have validated molecular resistance determinants (*8-13*).

Although the antimalarial drug development pipeline is strong, there are no current replacements for either the ART or partner drug component of ACTs that could be utilized in regions with high prevalence of drug resistant parasites. To address the lack of replacement options, alternative strategies using existing antimalarials are under evaluation. Among these are triple artemisinin-based combination therapies (TACTs), in which an additional partner drug is carefully selected and added to an existing ACT. The added antimalarial effect mediated by the third drug, ideally with a mechanism of action and resistance distinct from the other two compounds, would have the potential to slow down the emergence of resistance. In addition, TACTs could rescue ACTs in regions with partner-drug resistance, the main predictor of clinical treatment failure (*14*). Ideally, TACTs could also be combined in a manner that maximizes synergistic drug interactions or exerts counter-selective pressure on the parasite to thwart the acquisition of drug resistance.

Trials are currently ongoing to evaluate the safety, tolerability, and efficacy of TACTs. One recently completed trial evaluated DP+mefloquine and AL+amodiaquine, comparing the efficacy, safety, and tolerability to the ACT two drug combinations DP, ASM or AL (*15*). The study found both TACTs safe, tolerable, and as efficacious as the ACT comparative arms. Other combinations being evaluated include AL+amodiaquine and ASM+piperaquine (DeTACT-Africa, NCT03923725 and DeTACT-Asia, NCT03939104); ASP+atovaquone/proguanil and ASM+atovaquone/proguanil (NCT03726593); and AL+amodiaquine (NCT03355664). These studies will further elucidate the tolerability and safety of TACTs, and the feasibility of triple drug combinations for future treatment strategies to delay the emergence of drug resistant parasites.

Clinical evaluation of TACTs is essential, however such studies are expensive, time-intensive undertakings that require substantial resources. As funding for trials is also limited, TACTs with the greatest potential, regarding therapeutic efficacy, safety, and tolerability, should be prioritized. To facilitate the evaluation of potential combinations, we sought to develop a rapid, cost-effective process to prioritize combinations for clinical testing and identify genetic determinants that may modulate TACT efficacy and serve as genetic markers of decreased parasite susceptibility or drug resistance. Given the relative ease of *in vitro* high-throughput screening against *Plasmodium falciparum*, permutations of potential drug combinations can be rapidly interrogated to identify any synergistic or adverse pharmacodynamic interactions (altered antimalarial effects of the drugs in combination against the parasite, as determined by 72 hr proliferation responses). Similarly, culture-adapted parasites and genetically engineered lines that possess known or suspected genetic elements involved in drug resistance can be screened to test for negative pharmacogenetic interactions that might modulate the combination efficacy, thus threatening its longevity. This information could then be combined with single drug/ACT pharmacokinetic (and modeling), safety, and genetic epidemiology data to inform which TACTs to prioritize in clinical trials.

Given the complexity of the pharmacodynamic interaction between three compounds, this preclinical platform to screen TACTs uses a controlled *in vitro* assay to assess triple drug pharmacodynamic and pharmacogenetic interactions. Initial development and screening utilizing this process has demonstrated that an amplification of *plasmepsin II/III*, encoding two aspartic proteases involved in hemoglobin degradation, is associated with antagonism between two antimalarial drugs, PYR and piperaquine (PQP). This provides evidence for challenging the pairing of these antimalarials in a potential TACT, especially in Southeast Asia, where parasites with *plasmepsin II/III* amplifications are prevalent. Furthermore, antagonism between PYR and PQP was also noted when treatment was combined with antiviral protease inhibitors, supporting a broader implication for these drugs in combination therapies and highlighting the potential impact of co-administered medications for various comorbidities (*e*.*g*. HIV) on antimalarial therapy. This platform provides an efficient means for evaluating potential triple-drug combination regimens and the impact of concurrently administered medications on existing therapies, while also enabling further exploration of the chemical biology of *P. falciparum*.

## Results

### Optimization of triple drug screening platform

The experimental design employed strategies previously used to screen a diverse set of compounds in combination against *P. falciparum* lab isolates (*16*), with slight modifications to address neighbor-well effects and compound leaching (fig. S1). This method utilized an advanced acoustic dispense technology to first construct a 10-by 10-well matrix block within a 1536-well plate for each combination (Fig. 1), permitting a 9-point dose-titration for both compounds to be tested, along with the single drug activity. For triple drug combination screening, this 10×10 matrix block for the first two drugs was replicated 12 times in a 1536-well plate. The last block served as a DMSO control (*i*.*e*., no third drug present), allowing activity determination for each of the first two drugs alone and to assess the pairwise interaction. To each successive 10×10 matrix block, increasing concentrations of a third drug were added, with a total of 11 concentrations interrogated (with the same concentration of the third drug added to each well of the 10×10 matrix block).

**Fig. 1.**
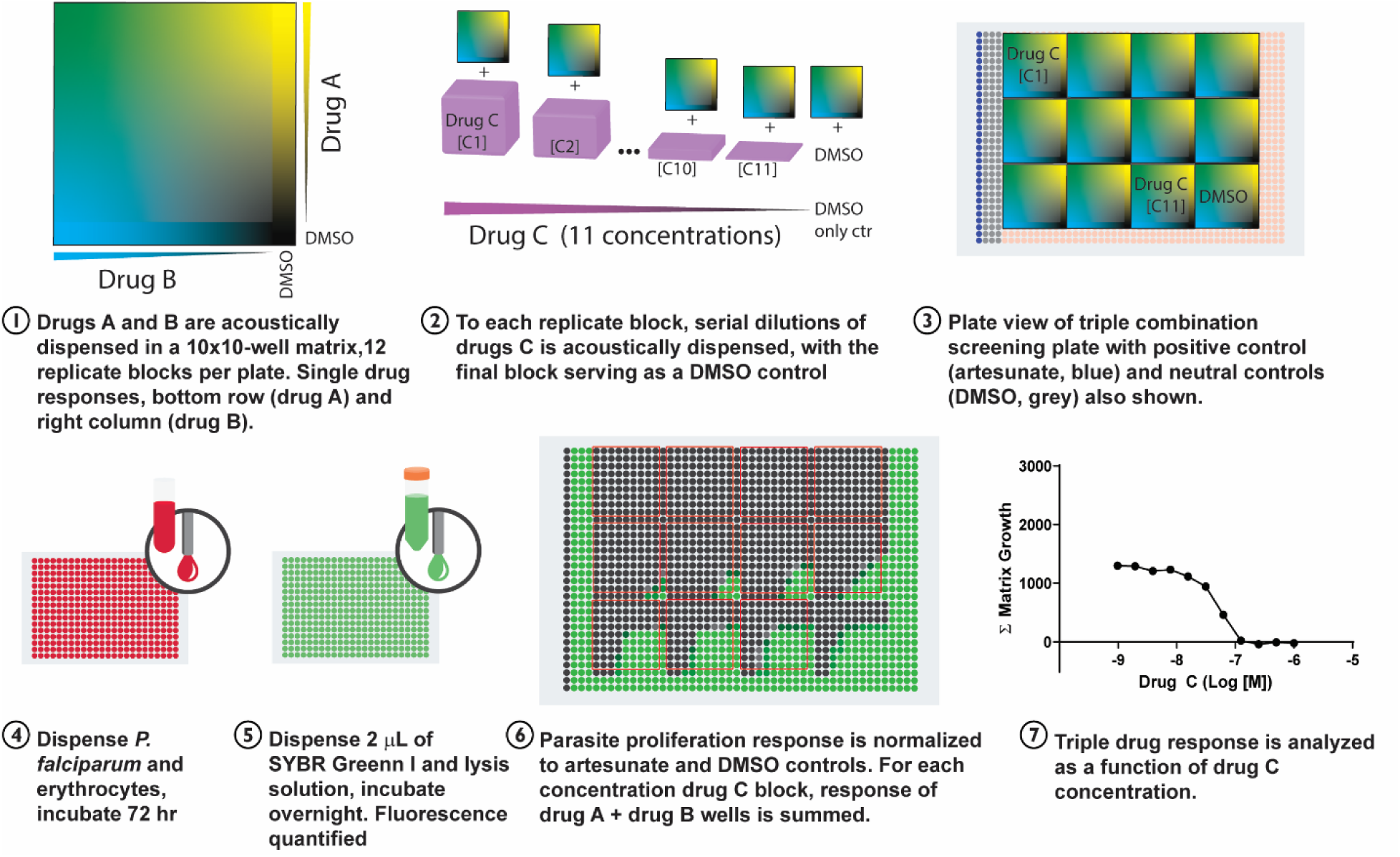
Quantitative high-throughput triple artemisinin combination screening methodology. Parasites were screened in 1536-well assay-ready plates. Briefly, drugs A and B (existing ACT combination drugs) were acoustically dispensed in a 10×10-well format with serial 2-fold dilutions. Drugs A and B were dispensed in the right-hand column and bottom row only, respectively, to allow assessment of the single drug response. This 10×10 matrix block was replicated across 12 independent blocks per plate, to which serial dilutions of drug C were added to each individual 10×10 matrix block. The last block served as a DMSO control. Parasites and erythrocytes were added in an 8 µL volume (2.5% hematocrit, 0.5% parasitemia) and plates were incubated for 72 hr. After incubation, SYBR Green I lysis solution was added. Fluorescence intensity of experimental wells was normalized to a 300 nM artesunate control column (1) and neutral DMSO control columns (2-4). For assessment of TACT response, the normalized response for each matrix block (81-wells in each block that received both drugs A and B, marked by red boxes) was summed and plotted as a function of drug C matrix block concentration.

Given the complexity of analyzing the activity and interactions of three independent drugs, all in dose titration, we sought an intuitive means for analysis and visualization. Raw fluorescent intensity values were internally normalized to an artesunate inhibitor (300 nM) and DMSO neutral controls, to calculate the normalized proliferation response. This normalization served to negate any inoculum or growth rate differences between assays. Normally, a drug’s potency is determined from a two-dimensional plot of the response (*e*.*g*. viability or proliferation) as a function of the inhibitor concentration with simplified metrics. For example, IC_50_, which is the concentration that inhibits 50% of the normalized growth response, IC_90_, and AUC (area under the dose-response curve) are determined by standardized growth/proliferation assays typically used for ranking drug potencies and comparative analysis. As the primary goal of this study was to evaluate the pharmacodynamic and pharmacogenetic interactions of potential TACTs and not specifically the dose-dependent interaction of the drugs, we initially compared the area under the TACT response derived from a volumetric assessment of the triple-drug response or by summing the first and second drug response and plotting a 2-dimensional representation of the response as a function of the third drug concentration for data analysis and visualization. Both methods yielded similar results, with the Σ matrix approach demonstrating easier computational and visual utility (Supplementary Methods; fig. S2). Due to the low drug C concentration in the 11^th^ dilution block (AQ, 0.4 nM; MFQ, 1 nM; LUM, 1 nM; PQP, 0.6 nM; and PYR 0.1 nM) the response was approximately equivalent to the DMSO (12^th^) control matrix block.

Although the evaluation of TACTs utilizes high-throughput screening methodology and light automation, it is limited as the assessment of the three drugs in dose-response requires one 1536-well plate per parasite line. As such, we focused our screening on culture-adapted field isolates to determine the efficacy against recent clinical isolates and assess if any genetic determinants modulate the drug responses. Shown in Fig. 2 are the Σ matrix responses for all 16 combinations (4 existing ACTs combined with 5 partner drugs) screened against 13 isolates, with inclusion of the PfKelch13 (mutations shown previously to modulate ART susceptibility) edited 967 isogenic lines, 967^K13C580Y^ and 967^K13R539T^ (*17*). As shown, there is variability in the TACT efficacy across the lines tested, and variability in the responses to individual TACTs for each line tested. As expected, decreased susceptibility, as measured by single drug response, to the individual second or third drug of the TACT modulated the Σ matrix response in lower concentration dilutions of the third drug. This is reflected in the correlation between the Σ matrix response and the single drug AC_50_ response, compound concentration that corresponds to 50% the normalized parasite growth inhibition (fig. S3).

**Fig. 2.**
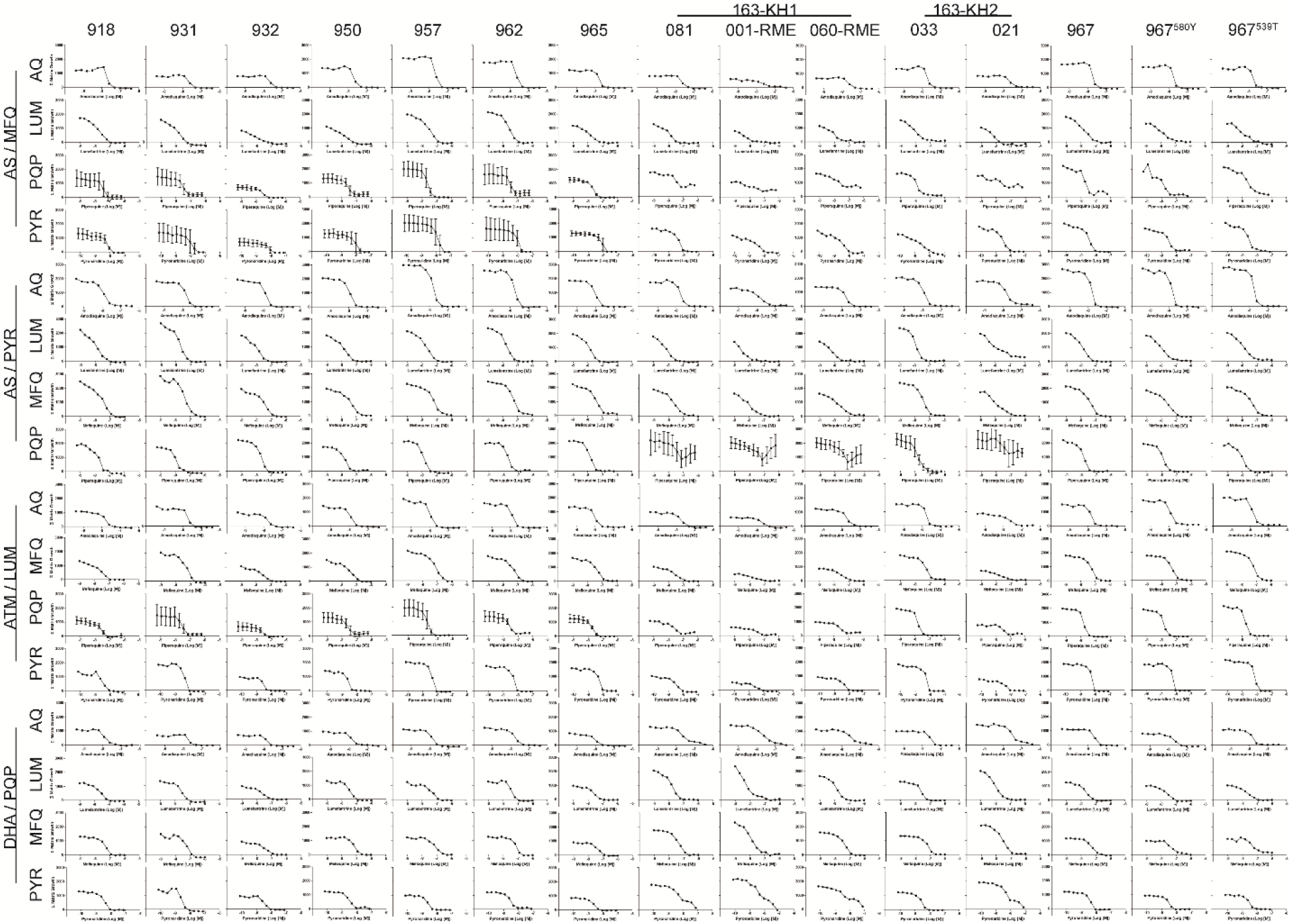
Overview of the Sum matrix responses. Sum (Σ) matrix responses of culture-adapted Cambodian field isolates and genetically modified lines after *in vitro* exposure to triple artemisinin combination therapies represented as 2-dimensional plots. The vertical axis displays the 16 combinations tested and the horizontal axis shows the 15 *Plasmodium falciparum* lines screened. Each individual graph shows the Σ matrix growth on the y-axis (summed normalized growth for the 81 wells with the first two compounds tested in 9-point serial 1:2 dilutions) as a function of the third antimalarial drug, screened in 11-point serial 1:2 dilutions. The adapted field-isolates 918-967 were collected from the Pursat province of Cambodia from 2009-2010 (*18*), and PfKelch13 edited isogenic mutants, 967^K13C580Y^ and 967^K13R539T^ (*17*). The 163-KH1 and 163-KH2 isolates were also collected in Cambodia between 2012-2013 from Pursat and Preah Vihear provinces, respectively (*19*). Select combinations against select parasite isolates were screened as independent replicates (n=3), shown as mean ± standard deviation (SD). AS, artesunate; ATM, artemether; DHA, dihydroartemisinin; MFQ, mefloquine; PYR, pyronaridine; LUM, lumefantrine; PQP, piperaquine; AQ, amodiaquine.

### Altered partner drug interaction between isolates

We noted that there was variance in the overall response to TACTs in the 2009-2010 field isolates (918, 931, 932, 950, 957, 962, 965), specifically to AS/MFQ/PQP, AS/MFQ/PYR and ATM/LUM/PQP. Isolate 932 was more susceptible to each of the combinations compared to the other isolates (fig. S4), with hyperAUCs (AUC of the entire TACT response curve) between 3,390 to 4,131 compared to a range between 5,836-15,517 for the other isolates (table S1). The increased potency of the TACTs was associated with higher susceptibility to both MFQ (40.2 ± 16 nM; mean AC_50_ ± SD) and LUM (21.4 ± 12 nM), the lowest AC_50_ against MFQ, and the second lowest against LUM (isolate 918 had an AC_50_ of 18.9 ± 9.7 nM against LUM; fig. S4, table S2). Interestingly, the altered TACT response was also associated with a differential partner drug-drug interaction, with subtle drug-drug interactions observed between PQP and either MFQ or LUM, with excess HSA (assessment of drug-drug interaction based on the Highest Single Agent model) values of 97.4 ± 53.3 and 34.7 ± 20.0 (mean ± SD) compared to isolate 962 (MFQ/PQP excess HSA 338.8 ± 137.8 and LUM 211.2 ± 117.0 or isolate 931 (MFQ/PQP excess HSA 466.8 ± 287.1 and LUM 364.2 ± 268.7) suggesting increased drug-drug antagonism in the latter isolates (fig. S4). The observed degree of pharmacodynamic interaction between these partner drugs was not entirely associated with the single drug susceptibility, with the highest degree of drug antagonism observed for isolates 931 and 950, both of which were intermediate in their susceptibilities to both MFQ and LUM, with isolate 950 having the highest AC_50_ to PQP (26.2 ± 9.2 nM, range 21.4-26.2 nM), suggesting a more complex pharmacodynamic-pharmacogenetic interaction than simply one driven by a drug resistance determinant (fig. S4).

### Cluster analysis of parasite responses at various concentration ranges

Cluster analysis of both combination and single agent responses for parasite lines was performed to elucidate similar phenotypic profiles. For each parasite line we quantified the TACT response, as a function of the area under the curve (AUC), for three ranges of the drug C concentration: low, middle and high concentration range. This gave rise to respective response-vectors. Additionally, an expanding-window based aggregation method was also applied to aggregate responses at different concentration ranges. As a baseline, the AC_50_ of the single agent response-vectors were determined for each drug for each of the individual isolates (table S2). The agreement between different clusterings was first characterized *via* pairwise adjusted-Rand indices (ARIs). The values for most of the comparisons are considered low, with a range of 0.11-0.83 (fig. S3). Indicating an overall low agreement of clusterings between parasite single agent and TACT responses. A potential explanation for the low agreement between the responses is that determining the resultant clusters requires cutting the dendrograms horizontally, which is a known difficulty associated with hierarchical clustering methods (*20-22*).

To address this, an alternative methodology to assess the response correlations was applied utilizing hierarchical dendrogram clustering and quantified by Cophenetic- and Baker-correlations (fig. S3). The highest correlation can be observed between clusterings produced by the expanding-window and the high and middle concentration response aggregation strategies (0.94 for both comparisons using Cophenetic, 0.94 and 0.93, respectively, using Baker-correlations). Expanding-window clustering is generally in agreement with the other clusterings (range 0.75-0.94 Cophenetic, 0.82-0.94 Baker). Excluding the expanding-window and single agent response-based clustering, the highest correlation is observed between the middle concentration range and other clusterings (range 0.78-0.86 Cophenetic, 0.81-0.87 Baker). Overall, the high agreement between different clustering methods suggests that the clustering of parasite response based on TACT or single agent phenotype is robust from the aspect of the concentration range at which the phenotypic response was investigated.

While the above measures can quantify the agreement of the clustering results, they do not elucidate the underlying topological similarities and dissimilarities of the dendrogram pairs. To address this limitation, we analyzed the dendrograms using a tanglegram approach that can provide insightful visualization and quantitative measure of agreement between the dendrograms. The resultant tanglegrams (fig. S3, table S3) demonstrate a high similarity by all clustering methods for the isolates 163-KH1-081, 163-KH1-001-RME, 163-KH1-060-RME and 163-KH2-021. For the other cell lines, the agreement between their clustering is less evident. The lowest degree of entanglement (0.13), highest degree of agreement, was observed for the single agent/TACT high concentration response vector clustering, with three larger, matching clusters present in both dendrograms. A similar, slightly less robust, agreement (0.14) is present in the single agent/TACT middle concentration range response vector clustering.

### Antagonism between piperaquine and pyronaridine is associated with plasmepsin II/III copy number variation

The relationship between single drug response and TACT response did not hold for all conditions tested. A striking difference in the Σ matrix response was noted for the TACT combinations containing both PYR and piperaquine (PQP) in the isolates collected from 2012-2013 (Fig. 3A and 3B). Most of these lines were able to proliferate to approximately 50% of the normalized response in the matrix block wells containing the highest concentration of PQP (600 nM) and PYR (75 nM; Supplementary Methods). Interestingly, this set of isolates (163-KH1 and 163-KH2 haplotype lines) was collected as part of an earlier study assessing DHA-PQP clinical efficacy in Cambodia (*19*), with those manifesting the altered Σ matrix response being the same lines associated with clinical treatment failure of DHA-PQP. Previous work demonstrated that these lines (163-KH1-081, 163-KH1-001-RME, 163-KH1-060-RME and 163-KH2-021) are associated with clinical treatment failure of the ACT, DHA-PQP, which is associated with a copy number amplification of *plasmepsin II/III* (table S4) (*11*). More recent work has demonstrated that the M343L mutation in the *P. falciparum* chloroquine resistance transporter (PfCRT), present in the 163-KH1-001-RME isolate can decrease the parasite susceptibility to PQP especially in the IC_90_ region of the concentration-response curve (table S4)(*12*). As suggested by the direction of the normalized SYBR Green I intensity results, pairwise combination matrix experiments demonstrated a strong antagonistic interaction between PYR and PQP in lines with amplified *plasmepsin II/III* (Fig. 3C); this interaction was not observed in the temporally related 163-KH2-033 isolate, which was successfully treated with DHA-PQP, but does not possess amplification of *plasmepsin II/III*. Furthermore, interaction for each of the *plasmepsin II/III* amplified lines demonstrated significant antagonism between PYR and PQP, as determined by the excess highest single agent (Excess HSA) interaction model (Fig. 3D). For each line that contains *plasmepsin II/III* amplifications we observed a significant amount of antagonism based on the HSA model ranging from 359.1 ± 209.6 for 163-KH2-021 to 427.1 ± 135.0 for 163-KH1-001-RME, whereas there was a trend towards a synergistic interaction in the 163-KH2-033 line, −212.1 ± 196.0, mean HSA model ± standard deviation (SD), n=9.

**Fig. 3.**
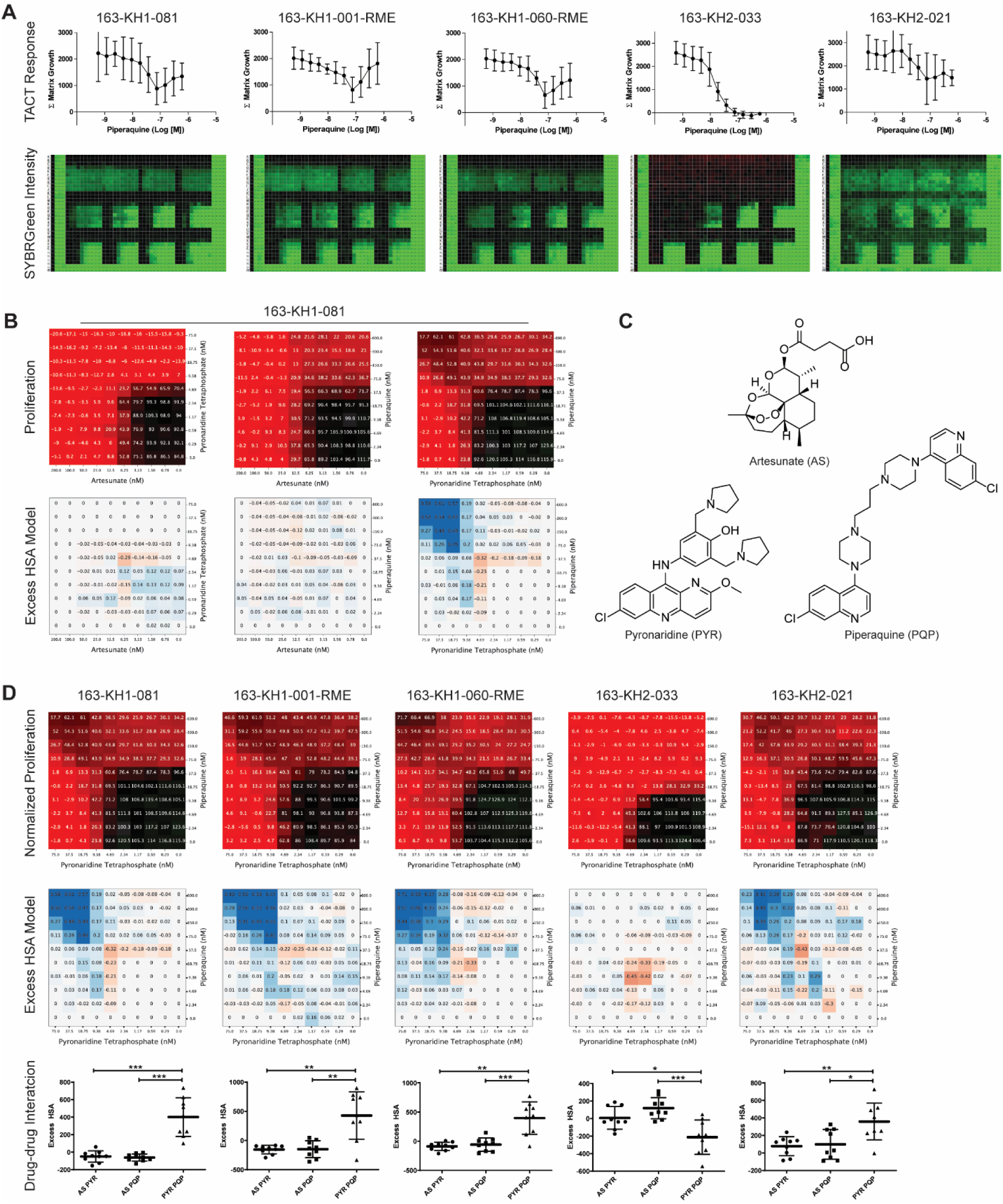
Pyronaridine/piperaquine antagonism associates with *plasmepsin II/III* copy number variation. **A)** Triple combination screening results for artesunate, pyronaridine (PYR) and piperaquine (PQP) (Σ (sum) matrix graphs display; mean ± SD; n=3). Representative normalized SYBRGreen I fluorescence intensity heatmap results for each line (green, high DNA content (parasite proliferation); black low DNA abundance (no parasite proliferation)). All lines were associated with clinical treatment failure of DHA-PQP, except 163-KH2-033. **B)** Two drug matrix assays demonstrate specific antagonism arising from the PYR/PQP interaction in the 163-KH1-081 line (black, parasite proliferation; red, no parasite proliferation. **C)** Compound structure for artesunate (AS), PYR and PQP. **D)** Interaction between PYR and PQP were evaluated for each of the 2012-2013 field isolates with the normalized response between PYR and PQP. The excess HSA model of drug-drug interaction visualization with blue representing antagonism, and red representing synergy. Quantified results from each pair-wise drug-drug comparison are shown below the HSA model (n=9; t-test assessment-*P* < 0.05, *; *P* < 0.01, **; *P* < 0.001, ***).

### Antagonism is not mediated by related 4-aminoquinoline molecules

As the aminoquinoline moiety is common to several clinically used antimalarial compounds (4-aminoquinolines-PQP, amodiaquine and chloroquine; 8-aminoquinolines-primaquine and tafenoquine) we sought to further interrogate if this moiety was mediating the antagonistic interaction with PYR. This was performed using pairwise matrix screening in combination with PYR. As shown in Fig. 4A, no other 4-aminoquinoline compound tested, amodiaquine, chloroquine or GSK369796, produced an antagonistic interaction with PYR. We also evaluated the interaction between PQP and quinacrine (Fig. 4B), a historical antimalarial that shares a tricyclic ring structure with PYR, and found a slight antagonistic interaction; however, this was not as pronounced as the PQP/PYR interaction. Further investigation exploring the mechanistic basis underlying the PQP/PYR antagonism in *plasmepsin II/III* amplified lines is ongoing.

**Fig. 4.**
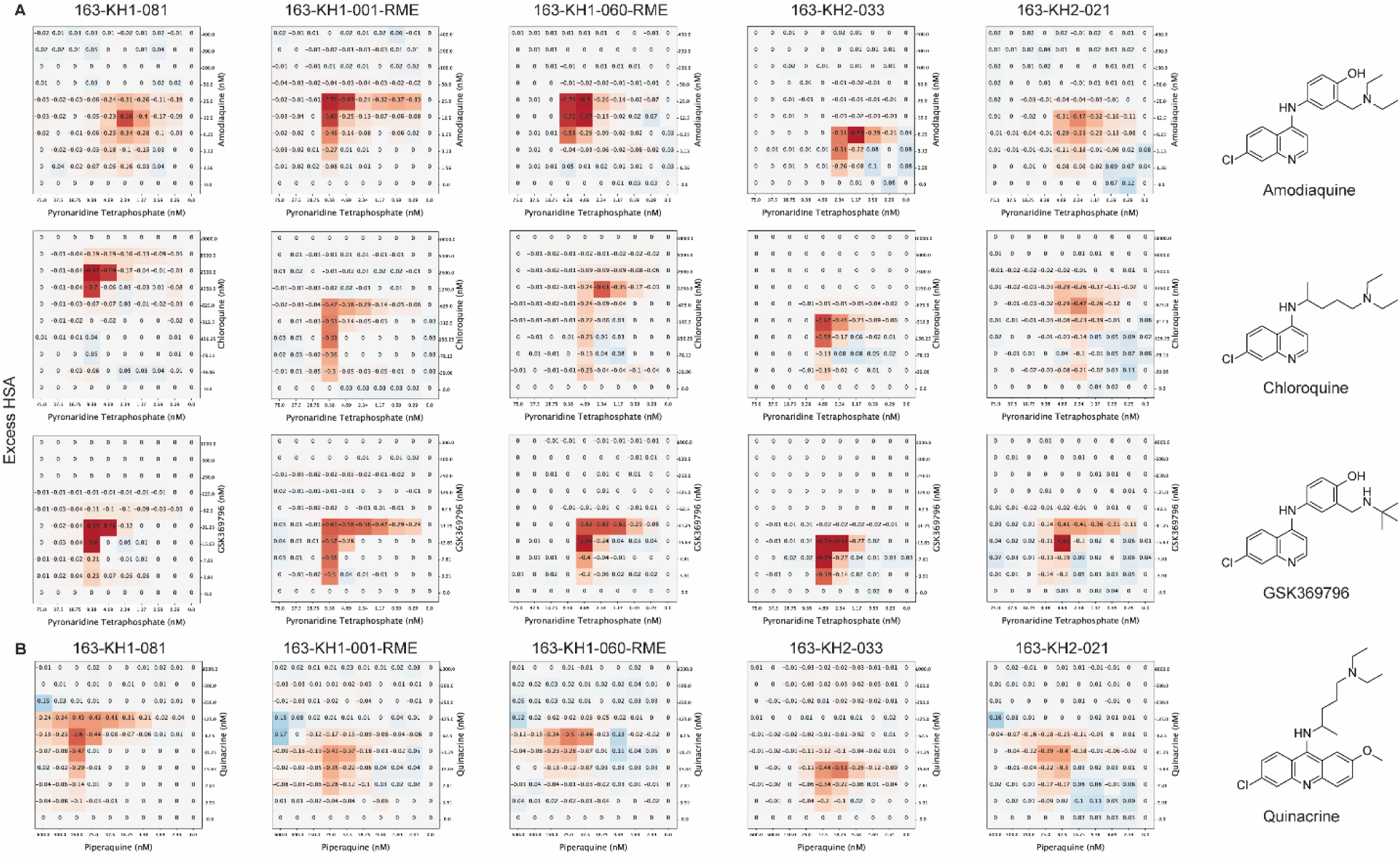
Antagonism between pyronaridine and piperaquine is not mediated solely by aminoquinoline moiety. **A)** To determine if other related antimalarials also demonstrated an antagonistic interaction we screened related compounds in a pairwise fashion. 4-aminoquinoline compounds, amodiaquine, chloroquine and GSK369796, did not demonstrate antagonism with pyronaridine in any of the lines tested. **B)** Quinacrine, which possesses a tricyclic structure similar to pyronaridine, demonstrated weak antagonism with piperaquine in lines that have amplified *plasmepsin II/III* copy number. Excess HSA drug interaction model color coding: white, no interaction; red, additive to synergy; blue, antagonism.

### Piperaquine/pyronaridine antagonism induced by aspartic protease inhibitors

Although the specific mechanism of action of PQP and PYR are unknown, evidence supports that both antimalarial drugs target the parasite digestive vacuole (DV), at least in part (*12, 23, 24*). As the observed antagonism was associated with parasites possessing amplified *plasmepsin II/III* copy numbers, both DV resident aspartic proteases involved in hemoglobin digestion, this suggested a means to interrogate the underlying molecular mechanism. A simple approach was used, in which we asked if a compound possessed the ability to reverse the antagonistic interaction between PYR and PQP. We focused our screen on protease inhibitors or molecules that may perturb DV function, such as protoporphyrin IX, the organic product released upon hemoglobin digestion. These compounds were screened against two lines that have amplified *plasmepsin II/III* copy numbers (163-KH1-081 and 163-KH1-001-RME) and 163-KH2-033, which has a single copy of both *plasmepsin II/III* and does not exhibit the PYR/PQP antagonism. None of the compounds screened demonstrated any reversal of the PYR/PQP antagonistic interaction (fig. S5). Surprisingly, we observed induced antagonism in the 163-KH2-033 line between PYR and PQP when treated with some HIV aspartic protease inhibitors (Fig. S5B). To investigate this further, we screened several adapted field isolates (none with amplified *plasmepsin II/III* copy numbers), and the lab line Dd2 to assess the induced antagonism between PQP/PYR and viral protease inhibitors (screened in 11-point dose response, similar to the TACT screening). Varying induction of antagonism was demonstrated for each line tested, with the HIV protease inhibitors ritonavir, saquinavir, atazanavir, indinavir and nelfinavir manifesting the largest degree of interaction (Fig. 5C and D). There is a degree of direct antagonism between PQP and certain PIs (Fig. 5C and D) that is not observed in comparison with PYR combinations. Of note, *P. falciparum* Dd2 is a drug-pressured line that originated from the Indochina III line isolated from a Laos refugee in August of 1980 (*25*). Thus, it has not been previously exposed to PYR or PQP, however of note, PQP was used as monotherapy in China in the late 1970s, but there is no reported usage in Laos (*23*).

**Fig. 5.**
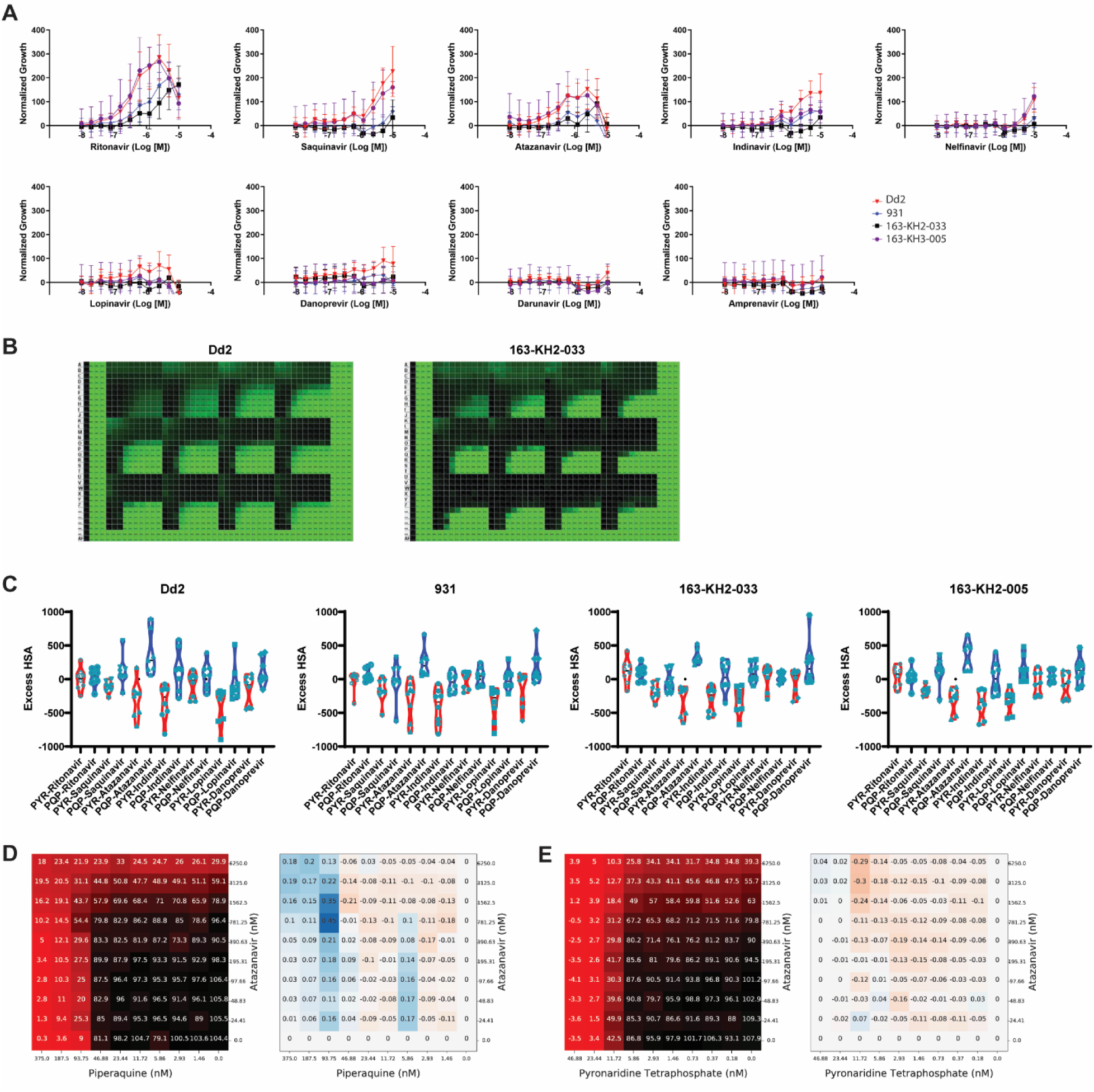
Viral protease inhibitors can induce antagonism between pyronaridine (PYR) and piperaquine (PQP) in lab and field isolates that do not have amplified *plasmepsin II/III* copy numbers. Based on the initial findings we profiled several viral protease inhibitors for mediating antagonism between PYR and PQP in dose-response. **A)** Normalized growth response of the top 16 PYR/PQP concentration wells treated with varying concentrations of protease inhibitors. Antagonism induction appears distinct from the single agent potency of the protease inhibitor (e.g. ritonavir and danoprevir both have AC_50_s against Dd2 of 3.9 µM). **B)** Normalized proliferation results of the ritonavir triple drug screen (PYR and PQP) for both *P. falciparum* Dd2 and 163-KH2-033 showing induction of antagonism in the higher ritonavir concentration matrix blocks in a dose-dependent manner (green, high DNA content (parasite proliferation); black low DNA abundance (no parasite proliferation)). **C)** Evaluation of the pharmacodynamic interaction of two-drug combinations, either PYR or PQP, and protease inhibitor interaction, shown is the mean ± SD, of 7-8 independent experiments. There does appear to be a more antagonistic interaction (larger, positive excess HSA value) between PQP and some viral protease inhibitors compared to PYR-protease interaction. Corresponding normalized proliferation and excess HSA responses for PQP-atazanavir (**D**) and PQP-atazanavir (**E**) for *P. falciparum* Dd2. The normalized proliferation responses are shown as: black, parasite proliferation; red, no parasite proliferation; and the excess HSA as blue, representing antagonism and red, representing synergy.

## Discussion

Multidrug combination therapies are now the standard of care for many diseases, including both communicable and non-communicable illnesses. However, the means to assess the optimal drug combination in an efficient and cost-effective manner remains a challenge for the development of such combination regimens, including the specific case of antimalarial combination therapies. Although some combination properties, such as modulation of pharmacokinetic properties mediated by host absorption and metabolism largely remain limited to *in vivo* models and clinical assessment, pharmacodynamic interactions and pharmacogenetic interactions can be assessed using *in vitro* assays, which enable the triaging and prioritization of combinations for more intensive or costly studies. Importantly, these high-throughput assays can quickly identify adverse multidrug pharmacodynamic interactions that may not be predictable based on single drug activities while also identifying genetic determinants that may modulate the effectiveness of the combinations which could then be utilized as genetic markers conferring a selection advantage to the combination pressure.

For antimalarial treatment, most endemic regions have transitioned to ACTs after the emergence and spread of drug-resistant parasites limited the utility of monotherapies (*23, 26*). However, growing resistance to partner drugs has resulted in failures of first-line ACTs, necessitating transitions to alternative partner drug combinations (*7*). For example, use of DHA-PQP has selected for parasites resistant to the PQP component that is associated with increased clinical treatment failure (*11, 27*). Although the antimalarial drug development pipeline is strong, due in large parts to efforts by the Medicines for Malaria Ventures (MMV), there is no ready replacement for the fast-acting artemisinin component and only limited clinically approved compounds that can be combined with an artemisinin component as an ACT. To address this immediate need, one option is the addition of another partner drug to an existing ACT, creating a TACT with improved overall combination efficacy that can limit the evolution and spread of drug resistant parasites.

There are currently five ACTs listed on the WHO Model list of Essential Medicines (*28*), yielding 10 possible TACT combinations, not factoring for the inclusion of an antimalarial not currently part of an ACT, or late stage clinical candidates. Evaluation of all possible TACTs through *in vivo* models or in a clinical setting would require intensive resources. As such, we developed and evaluated a high-throughput quantitative process for evaluation of three-drug antimalarial combinations to inform on the prioritization of combinations for further clinical evaluation. As the emergence of parasites with reduced susceptibility to both components of the ACT DHA-PQP have arisen in the Greater Mekong Subregion, we used adapted field isolates from this region to both evaluate the TACTs in the context of multidrug resistant isolates and also elucidate pharmacogenetic interactions that may exist against these circulating parasites.

Initial clinical evaluation of TACTs for the treatment of uncomplicated *P. falciparum* malaria was assessed across four countries in Southeast Asia. Work by van der Pluijm *et al*. has recently evaluated two TACTs: DHA/PQP+MFQ and ATM/LUM+AQ compared to the existing ACTs DHA/PQP and ATM/LUM, respectively (*15*). Regarding adverse events, there was a slight increase in vomiting in the TACT treatment arms compared to the ACT treatments, as well as a non-clinically relevant extension of the QTc interval with ATM/LUM+AQ. However, the study notably demonstrated that the incidence of serious adverse events was similar for TACTs and ACTS (*15*). Overall, this study supports the efficacy, safety, and tolerability of TACTs. Addition of MFQ to the existing DHA/PQP ACT resulted in an increased clinical efficacy of the treatment, at all sites with 97% efficacy of the TACT versus 60% for the ACT (in a setting of previous DHA-PQP treatment failures). Additionally, ATM/LUM and ATM/LUM+AQ had similar efficacy across all sites (97% versus 98%, respectively) (*15*). The increased efficacy of DHA/PQP+MFQ that was observed in Southeast Asia with a high prevalence of parasites resistant to PQP (leading to a decreased efficacy of the ACT DHA/PQP), supports the application of TACTs for thwarting drug resistant parasites. By demonstrating that TACTs remain potent against individual drug resistant isolates, these findings support the hypothesis that TACTs would decrease the emergence of drug resistant parasite populations by limiting their spread and thereby increasing the longevity of the combined therapeutic regimens.

Overall, our findings demonstrate a utility for screening TACT combinations *in vitro*. Based on the limited combinations and parasites screened as part of these initial efforts, we were able to identify combinations of PYR and PQP that have an antagonistic interactions in parasite lines with amplified *plasmepsin II/III* copy number variations (CNVs), a highly prevalent genotype in the Greater Mekong Subregion (*11, 29, 30*). These isolates were collected as part of a DHA-PQP efficacy study that demonstrated decreased clinical efficacy of the ACT (*19*). Since they were recently adapted field isolates and reflective of circulating genotypes, they were included in this study. Although there is an association between *plasmepsin II/III* CNV and decreased PQP susceptibility, controlled experiments fail to demonstrate, at least in the *P. falciparum* 3D7 genetic background, that overexpression of either *plasmepsin II* or *III*, directly affect the ability of this CNV to modulate PQP susceptibility (*31*). Indeed, current genetic epidemiological evidence supports a role for mutations in *pfcrt* in conferring resistance to PQP (*12, 30, 32-34*). Studies by Silva *et al*. have also suggested that the *plasmepsin II/III* amplifications and *pfmdr1* deamplifications have created a genetic background that favors the accumulation of *pfcrt* mutations (*35*). Further investigation to validate and elucidate the molecular mechanism mediating the induction of an antagonistic interaction between PQP and PYR in the context of *plasmepsin II/III* CNV will be required, which may also aid in understanding the specific mechanism of action of these two antimalarial compounds.

Association of the antagonistic phenotype between PQP and PYR in field isolates possessing amplified *plasmepsin II/III* suggested a potential molecular mechanism. Plasmepsin II and III are aspartic proteases that localize to the parasite DV wherein hemoglobin degradation occurs (*36*). The DV is also a proposed site of action for both PQP and PYR (*37*). As such, we hypothesized that increased expression of *plasmepsin II*/*III* yielded increased protease activity, thus altering the DV environment in a manner that facilitated an antagonistic interaction between PQP and PYR. However, screening a limited set of protease inhibitors and other molecules that could alter the DV function supported an opposite conclusion. Shown in Fig. 5 and fig. S5, treatment with HIV aspartic protease inhibitors had no significant effect on the antagonism of PQP and PYR in parasites with amplified *plasmepsin II* and *III*, but induced an antagonistic interaction in both lab and field isolates lacking the CNV. Pairwise HIV protease inhibitor (PI) evaluation demonstrated a more antagonistic interaction with PQP compared to PYR (Fig. 5). However, this does not seem to wholly explain the triple drug interactions observed, as the concentration of PQP and PYR individually were able to abrogate parasite growth, yet addition of protease inhibitor increased the viability of the PQP/PYR/PI treated wells to ∼50% of the neutral control. This suggests a complex pharmacodynamic interaction, possibly through modulation of the DV environment, affecting the mode of action of the antimalarials or altering parasite metabolism and thereby permitting parasite proliferation.

Developing TACTs presents a challenge regarding both unknown drug pharmacodynamic interactions as well as optimizing pharmacokinetic properties while minimizing toxicity. Utilization of *in vitro* and *in vivo* models, together, can facilitate the informed selection of combinations to put forward for evaluation in clinical trials. High-throughput *in vitro* assessment of combination potency enables initial triaging of combinations to evaluate drug interactions and impact of drug resistance determinants before employing more labor intensive and costly means (*i*.*e*., *in vivo* models and clinical assessment). Identification of combinations using this *in vitro* platform with negative attributes (*i*.*e*., strong antagonism) would likely be predictive and can be used to deprioritize combinations for downstream testing. Additionally, identification of genetic determinants that modulate TACT potency could also be useful for prioritizing genetic markers for molecular epidemiological follow-up. Although *in vitro* methods are able to capture pharmacodynamic and pharmacogenetic interactions, more labor-intensive *in vivo* methods or clinical assessment will be required to evaluate the TACT pharmacokinetic and toxicity interactions. Importantly, *in vitro* phenotypic screening (such as ability to inhibit parasite proliferation) for infectious diseases has high predictive validity of clinical therapeutic response (*38, 39*). Although initial TACT clinical evaluations found no significant risk of altered QTc interval prolongation (*15*), there remains the potential increased cardiotoxic risk arising from the use of TACTs containing quinoline antimalarials. As such, further monitoring of potential cardiotoxicities of TACTs, with or without co-administered therapies (*e*.*g*., ritonavir) that may contribute to QTc interval prolongation, should be continued (*40*).

The concentration ranges of drugs used in this study was optimized based on the *in vitro* potencies (AC_50_) for all parasite lines and does not specifically recapitulate the physiologic concentrations or kinetics of drug exposure *in vivo*. In a population pharmacokinetic analysis study, Hoglund *et al*. found that PQP had a median C_max_ of 248 ng/ml (24.3-1,170 ng/ml, minimum-maximum) and a half-life of 22.5 days (9.15-52.3) (*41*). This would equate to an approximate C_max_ of 463 nM (range 45-2,187 nM), which compares with our *in vitro* concentration range of 0.6-600 nM for PQP. PYR was shown to have a median C_max_ of 341 ng/ml (226-571 ng/ml, minimum-maximum) with a half-life of 16.5 days (10.9-26.5), corresponding to an approximate concentration of 658 nM (436-1,102 nM) (*42*). This reflects a higher achieved concentration *in vivo* compared to the range used in these *in vitro* studies (0.1-75 nM). Although given the variabilities in the *in vivo* achievable C_max_ and shorter half-life of PYR, it is likely that similar ratios of the drugs are achievable during the elimination phase. Regardless, these high-throughput assessments of antimalarial combinations are meant to provide insight into the complex pharmacodynamic interactions that would then be explored with more labor-intensive methodologies or evaluated clinically (*43*).

This methodology can also be utilized to identify pharmacodynamic interactions between antimalarial combinations and other medications that may be taken during treatment for malaria. Our finding that HIV PIs can modulate antimalarial drug-drug interactions highlights an understudied aspect of antimalarial therapy, the impact of co-administered medications. Importantly, the concentrations of HIV PIs, some of which modulated efficacy in a dose-dependent manner (e.g. ritonavir and atazanavir, see Fig. 5A) were tested at levels which approximate clinically relevant concentrations, or concentrations that would be achieved in patients on standard antiretroviral dosing regimens (*44*). This is particularly important when considering that malaria occurs in many parts of the world where HIV prevalence remains high (*1, 45*). Although HIV PIs are not first line treatments in adults, they are second-line therapies in younger patients (adolescents and younger children as well as infants) and remain a mainstay of primary therapy (*46, 47*). With millions of people on or needing access to antiretroviral therapy, understanding the implications of pharmacodynamic interplay between HIV PIs and antimalarials remains integral. Indeed, the HIV PIs which had the most pronounced effect in our hands included ritonavir, which is a boosting-second agent universally used with any of the primary PIs, if HIV PI-based antiretroviral therapy is going to be used (e.g., ritonavir-boosted lopinavir, atazanavir, and darunavir are all commonly recommended regimens for children on HIV PI-based ART). Although use of TACTs has been supported by initial large-scale clinical trials (*14, 15*), appreciating that there may be potential negative interaction between TACTs and PIs is important in clinical HIV and malaria management strategies. This of course could be extrapolated to other therapies for treatment or other chronic diseases which may modulate the efficacy of antimalarials.

In addition to addressing the immediate need for prioritization of potential TACT combinations for clinical evaluation, this platform can be employed for evaluation of candidate antimalarial combinations during preclinical and early clinical development. Deployment of TACTs with existing partner drugs is a stopgap measure, as many of the existing partner drugs have known genetic determinants that can decrease drug efficacy. Ideally, novel triple-drug combinations without pre-existing drug resistance would be developed and deployed to maximize the lifespan of the combination and protect each component from the emergence of drug resistance (*48, 49*). *In vitro* assessment of pharmacodynamic and pharmacogenetic interaction would complement existing tools for the development of antimalarial combinations, including the Medicines for Malaria Venture (MMV) *in silico* Combo tool, development of combinations based on single agent *in vitro* and *in vivo* models, pharmacokinetic properties, safety data, drug resistance potential and chemical characteristics as well as evaluation of combinations *in vivo* in humanized mouse models and human challenge models (*50-52*). This platform could also be utilized for the evaluation of sequential ACT therapy, another proposed modification to existing ACT therapy to improve clinical efficacy and reduce the emergence of drug resistant parasites. In such protocols two different approved ACTs would be administered sequentially over 6 days (two three-day treatments), ideally with partner drugs with validated mechanisms of action that exert opposing pressure on genetic determinants that modulate drug susceptibility (*53*). During the second half of treatment, there would be remaining exposure of the initial ACT partner drug, thereby establishing a three-day timeframe of triple-drug exposure.

In conclusion, the simplified *in vitro* assays and associated analysis tools presented here provide a scalable platform to explore the dimensionality of triple drug combination therapies against *Plasmodium* parasites. Potential combinations can be rapidly and efficiently characterized *in vitro* providing valuable insight for the prioritization of compounds for more intensive preclinical and clinical evaluation. Furthermore, this work demonstrates the utility of profiling diverse genetic *P. falciparum* isolates for the identification of genetic determinants that impact the efficacy or modulate the pharmacodynamic interaction of combinations as well as screening potential co-administered medications for adverse interactions with antimalarial therapy. The establishment and greater utilization of this methodology facilitates early evaluation of pharmacodynamic and pharmacogenetic interactions for multidrug therapies, relevant for antimalarial development as well as other infectious and non-infectious diseases.

## Materials and Methods

### Parasite culture and quantitative high-throughput screening (qHTS)

*Plasmodium falciparum* parasite lines have been previously described (*18, 19, 54-57*) were cultured *in vitro* using standard conditions (*58*). Briefly, parasites were maintained in 2% human O+ erythrocytes (Interstate Blood Bank, Memphis, TN) in RPMI-1640 medium (Life Technologies, Grand Island, NY) supplemented with 0.5% Albumax II (Life Technologies), 24 mmol/L sodium bicarbonate, and 10 µg/ml gentamicin. Tissue culture flasks and assay plates were incubated at 37°C under a gas mixture of 5% CO_2_, 5% O_2_ and 90% N_2_. Methods for the SYBR Green-I qHTS, calculation of AC_50_ (concentration able to inhibit 50% of maximum parasite response) and definition of curve classes have been described previously (*59-62*). All HTS assays were read at 72?hr. Percent response values represent relative growth as judged by SYBR Green-I fluorescence intensity values normalized to controls.

### Combination assay

Two-drug combination and triple-drug combination assays were conducted as above for SYBR Green-I qHTS (table S5). Compounds were dissolved to 10 mM in 100% DMSO and stored in Matrix 2Dbarcoded tubes. Prior to screening drug combinations, the single-drug activities of the test compounds were evaluated against each of the lines to be screened. Compound dose ranges were optimized to encompass the AC_50_ value against the plurality of lines screened; for dihydroartemisinin, artemether and artesunate these ranged from 200-0.78 nM; amodiaquine ranged from 400-0.4 nM; mefloquine and lumefantrine ranged from 1000-1 nM; piperaquine ranged from 600-0.6 nM; and pyronaridine ranged from 75-0.1 nM. All assays were screened using the same concentration range and serial 1:2 dilution scheme, partner drug dilution in the replicated 10×10 matrix block was performed from the same top concentration but was 9 serial 1:2 dilutions. Compound source plate for acoustic ejection transfer to assay plates is described in Supplemental Methods. SYBR Green I fluorescence values were used to calculate a normalized response for each plate tested, based on the inhibitor control (artesunate 300 nM) and DMSO (neutral solvent) values. Utilizing the normalized response values, a Σ matrix response, based on the simple sum (SimSum) method, described below, was calculated to each 10×10 matrix block. This was calculated by summation of the 81 wells to which both compound A and B were added (Fig. 1). The Σ matrix response was then graphed as the SimSum calculation for each block plotted as a function of the third drug concentration.

### Sum matrix calculation and analysis methodology

As a general simplifying proxy of both potency and efficacy, the concept of calculating a “volume under the surface” (VUS) for specific dose-response grid “slices” was used. This calculation, analogous to area under the curve for single-agent dose responses, attempts to capture the total responsive effect seen across all doses in a 2D dose-response surface and can therefore be used to analyze how varying the concentration of a third agent affects the combined responses of the other two agents. The VUS is meant to be calculated in log concentration space and is used for relative comparisons within the same concentration structure. In the case of uniformly applied serial dilutions, concentration spacing can be considered unit-length (of note a 1:2 dilution series was used for all TACT assays, with fixed concentration used for each compound for all assays). To calculate the VUS, two distinct conceptual models for a surface were considered: simple sum, (“SimSum”) and “Mesh”, described in Supplemental Methods.

The SimSum volume model is the simpler of the two models. The surface of the dose-response matrix is constructed simply by extruding equally sized squares from their 2-dimensional log concentration grid locations. The advantage to this is model is that, with unit-length widths of extruded squares, the volume under the surface can be calculated as the SimSum of concentrations:

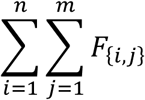

F is the two-dimensional dose-response matrix for agents A and B, with elements at i=0 having zero centration of agent A, elements at j=0 having zero concentration of agent B.

### Assessment of drug interactions

Plated wells were deconvoluted into individual response matrices and their combination behavior (additivity, synergy, antagonism) was characterized using a variety of metrics based on the Loewe, Bliss, and Gaddum models, as previously described, results from all metrics are available at https://tripod.nih.gov/matrix-client/ (*16*). Briefly, the Highest Single Agent (HSA) approach (also referred to as the Gaddum’s noninteraction or cooperative effect) simply reflects the fact that the resulting effect of a drug combination (*E*_*AB*_) is greater than the effects produced by its individual components (*E*_*A*_ and *E*_*B*_) (*63, 64*). Although this methodology has limitations assessing some occurrences of synergy, a positive result obtained with the HSA approach indicates an antagonistic drug-drug interaction compared to the single drugs considered alone.

Heatmaps for parasite AC_50_ response and TACT AUC response were generated using the Morpheus web-based visualization tool (https://software.broadinstitute.org/morpheus).

### Statistical analyses

Statistical significance was determined using a t test (unpaired, two-tailed), performed using GraphPad Prism (GraphPad Software Incorporated).

## Supplementary Materials

Supplementary Methods

Fig. S1. 1536-well assay optimization

Fig. S2. TACT calculation analysis methodology

Fig. S3. AC_50_ - AUC Relationship Analysis

Fig. S4. Drug interaction in 2009-2010 isolates

Fig. S5. Screen for chemical inhibitors of PYR/PQP antagonism

Table S1. Area under the triple artemisinin combination response curve

Table S2. Mean AC_50_ values for single agent screening against *Plasmodium falciparum* lines tested

Table S3. Entanglement of dendrograms generated by different clustering methods

Table S4. Parasite genotypes

Table S5. *Plasmodium falciparum* SYBRGreen assay 1,536-well protocol

## Acknowledgments

We would like to thank all the participants and families that participated in the previous studies from which the parasite isolates were collected (NCT00341003 and NCT01736319). In addition, we would also like to thank Craig Thomas (NCATS) for useful discussions and feedback. Genome sequencing was performed by the Wellcome Sanger Institute (WSI), and sequencing data was processed by the MalariaGEN Resource Centre. We thank the staff of the WSI Sample Logistics, Sequencing, and Informatics facilities for their contribution.

## Author contributions

MRA, CA, RvdP, RTE designed the experiment; MRA, ZI, JMS, RTE performed experiments; MRA, LC, GZ-K, CA, TP, CVH, JMS, RvdP, TEW, AS, RTE analyzed data; MRA, OM performed genetic analysis; SS isolated/culture adapted parasites; AMD, TEW, AS, RTE supervised experiments; MRA, CA, RTE wrote the manuscript with input from all authors.

## Data and materials availability

All normalized combination and triple drug proliferation response results along with drug-drug interaction calculations (Loewe, Bliss, and Gaddum models) are available at (https://tripod.nih.gov/matrix-client/?project=2321). All clustering correlation related data analysis and R scripts are available at GitHub (https://github.com/ncats/tact.git).

