## Supplement Materials for "Modulation of triple artemisinin-based combination therapy pharmacodynamics by *Plasmodium falciparum* genotype"

#### Supplementary Materials

#### Supplementary Methods

*Dose Matrix Block Construction Using Acoustic Droplet Ejection*

An in-house software package, Matrix Script Plate Generator (MSPG), was used to create all necessary files for generation of compound source plates, acoustic dispensing scripts and plate maps for each TACT (*65*). To generate the compound source plate, a Perkin-Elmer Janus Automated Workstation with disposable Varispan tips was used to transfer compounds from 1.4 mL Matrix 2D barcode tube (sample tube) to the individual wells in columns 1 and 11 of Echo Qualified 384-Well Polypropylene Microplate 2.0** (Labcyte, San Jose, CA). Drug A, B and C for each TACT were plated multiple times to provide enough compound volume for generation of assay plates to screen each *P. falciparum* line using a single drug source plate. The identities and positions of each sample were recorded in an accompanying Excel spreadsheet. DMSO was dispensed to wells in columns 2-10 and 12-22 of Echo qualified 384-Well Source plate containing compounds using a Matrix Wellmate (ThermoFisher Scientific, Waltham, MA). The plates were briefly centrifuged for 2 min at 100 × *g* in a benchtop centrifuge prior to serial dilution on Beckman Coulter Biomek FX (Brea, CA) which simultaneously performed serial dilutions for 9 to 11 dilution points from columns 1 and 11. The last column of every dilution series (10 and 22) remained as DMSO only. In addition, artesunate and DMSO solvent control were also transferred to columns 23 and 24 for dispense, serving as the positive (inhibitor) and neutral (solvent) controls. Customized (http://tripod.nih.gov/matrix/coma/) generated input files for the Labcyte acoustic dispenser (Labcyte, San Jose, CA) from a comma-separated value file that contained the unique plate and well pairings for each of the initial compound A and B matrix blocks, compound C addition was imputed separately using the Labcyte Reformatter software. Plate maps were simultaneously generated by the software to enable subsequent data normalization and analysis.

To achieve customizable dose–response matrix assays for three distinct compounds, we employed an Access Laboratory Workstation (Labcyte). Organization of each assay plate in true physical space resulted in 12 10×10 replica matrices within one 1,536-well plate, with one matrix serving as a DMSO control (drug A and B matrix alone) and the remainder receiving increasing concentrations of drug C. A single plate can be created in ~9 min, affording full plating of each

TACT combination of 16 plates in approx. 1.5 hr. The Labcyte Echo constructed the matrix blocks by dispensing 10 nL of each of the 9 wells associated with drug A down each column of every 10×10 block in 1536-well. Then, 10 nL of each of the 9 wells of each drug B dilution was dispensed across the 9 rows of their respective matrix block from left to right. Each plate was sealed with a peel-able aluminum seal, remaining covered until the initiation of the biological assay. Assay ready plates were, if not used immediately, frozen at -80˚C.

*P. falciparum parasites*, 0.5% parasitemia were dispensed in an 8 μL along with human erythrocytes (2.5% hematocrit final) were dispensed using a Multidrop Combi (ThermoFisher; table S5). Assay plates were then incubated for 72 h at 37°C in an atmosphere of 90% $N_2$/5% $O_2$/5% $CO_2$ with 95% relative humidity. All assays were read at 72 h using SYBRGreen-I based quantification of parasite proliferation, as previously described (*59-62*). Percent response values represent relative growth as compared to SYBRGreen-I fluorescence intensity values normalized to controls (artesunate 300 nM final concentration as positive control and DMSO neutral control), using formula: normalized growth % = (X – positive Ctrl) / (neutral Ctrl – positive Ctrl) × 100%.

*Mesh matrix calculation and analysis methodology*

The Mesh volume model is constructed by forming a vertex for each non-zero concentration pair in log-space and raising or lowering the vertex point in the third dimension relative to the measured response. The mesh is the surface formed by connecting adjacent neighboring vertices together. This VUS attempts to account for the "internal" surface, and ultimately weighs the internal measurements concentrations of the grid more highly than the edges. The VUS for the Mesh can be approximated by considering the center point of each 4-vertex distorted plane and taking the simple sum.

$$\frac{1}{4}\sum_{i=1}^{n-1}\sum_{j=1}^{m-1} \mathrm{F}_{\{i,j\}} + \mathrm{F}_{\{i+1,j\}} + \mathrm{F}_{\{i,j+1\}} + \mathrm{F}_{\{i+1,j+1\}}$$

Both VUS methods show utility as simplifying single-value potency and efficacy proxies facilitating the relative interpretation and visualization of the triple-drug response. As the concentration of third agents are varied in 3 agent screens, the VUS can inform some broad elements of synergy, antagonism, and concentration sensitivity. The choice of SimSum or Mesh as VUS scheme was not found to meaningfully alter the visual interpretation of results. Furthermore, a sample of 100 representative VUS calculations for both SimSum and Mesh found that the two results were very highly correlated (R-squared = 0.993, $P$<0.0001). Due to its relative simplicity, therefore, the SimSum method of summing the individual responses was specifically selected to serve as a major calculated feature.

*Cluster analysis of parasites responses at various concentration ranges*

Area-under-the-curve (AUC) was computed for various concentration-ranges of the dose-response curves associated with a specific cell-line and a triple-drug combination applied at various concentration-ratios. This gave rise to real-value vectors, i.e. response vectors, that characterize the response of a cell line with respect to a drug combination. Each dimension of a response vector represents a triple-drug combination. Note that a response vector can also be constructed from $AC_{50}$ values in an analogous manner to using AUC values. Such a response-vector was also computed for cell-lines.

For performing principal-component analysis of response vectors, the response-vectors were normalized by zero-centering and scaling the data along the dimensions (*66*). The normalized response-vectors were subject to principal-component analysis (PCA) (*66, 67*). The number of principal components (PCs) to withhold was determined with the help of generating scree-plots (*68*), and the input data was projected to these PCs. Transforming the response-vectors with PCA analysis allowed for computing the distance-matrix between pairs of vectors by employing the Euclidean distance metric (*69*).

Cluster analysis comparison of response-vectors allowed for identification of cell lines that respond similarly to different drug combinations. This analysis was performed with the help of cluster analysis as follows. Clustering was performed by utilizing Ward's-method which is an agglomerative clustering method and produces a so-called dendrogram (*70*). The resultant clusters can be extracted by determining the level of hierarchy at which the dendrogram needs to be cut. To this end, the Kelley-index of each dendrogram was determined, and the clusters were identified accordingly (*71, 72*).

The responses of cell lines associated with various concentration ranges and with $AC_{50}$ was subject to cluster analysis. We investigated the high, middle and low concentration ranges. Furthermore, we utilized an expanding-window based aggregation of AUCs. The expanding-window was calculated as a factorial of the SimSum VUS (SimSum VUS of block 11 (lowest concentration drug C), block 11 + block 10, 11 + 10 + 9, *etc.*). Therefore, for a given cell line five types of response vectors were computed giving rise to five different clusterings. Each clustering grasps a different aspect of cell response based on the underlying concentration range or $AC_{50}$ that was investigated. We sought to characterize the agreement of the five types of clusterings. To this end, we utilized three approaches.

Adjusted Rand-index. Agreement of two clustering can be quantified by investigating if pairs of objects were assigned to the same or different cluster in the two clustering outcomes. This information is quantified by the adjusted Rand-index (ARI) ranging from 0 to 1, the latter indicating perfect agreement (*73-77*).

Correlation of Dendrograms. Agreement of clusterings can be expressed based on the heights at which pairs of response-vectors are joined. The Cophenetic correlation first expresses the correlation between the pairwise Euclidean distances of response-vectors and the height at which they are first connected (*78, 79*). This correlation matrix is computed for each clustering. Then, the correlation of two clustering is defined as the correlation between the respective matrices, pairwise. A related method, i.e. the Baker-correlation (*80-82*), analyzes pairs of response vectors and the maximal height, i.e. level, at which the dendrogram can be cut so that the two vectors will still belong to the same cluster. For each pair of response-vector this level is determined for both clusterings. After matching the response-vector pairs in both clusterings, the Spearman-correlation between the levels is computed (*82*), giving rise to the Baker-correlation of two clusterings (*79*). The pairwise-correlations of clusterings are visualized with the help of correlation plot (*83*).

Tanglegram. Tanglegrams are used to compare the relations of a pair of dendrograms in a transparent manner. The positions of leaves are connected in both dendrograms. In the case of a perfect agreement the resultant tanglegram will connect all corresponding leaf-pairs with a straight line. Any deviation from the perfect agreement would introduce angled-connections between leaf-pairs, hence increasing the entanglement. The value of entanglement is influenced by the choice of distance-norm, that controls the magnitude of the penalty for angled-lines. In this study, we applied the L2 distance-norm. In the resultant tanglegram, identical subtrees are connected with colored lines. Also, dotted branches indicate separation of leaves that end-up in different sub-trees in the two dendrograms.

All clustering correlation related data analysis and visualization was conducted in R language (*66*) using various libraries; dplyr (*84*), reshape2 (*85*), ggplot (*86*), cluster (*69*), dendextend (*79*), mclust (*77*), corrplot (*83*), maptree (*71*) and stringr (*87*). Scripts have been deposited to GitHub (https://github.com/ncats/tact.git).

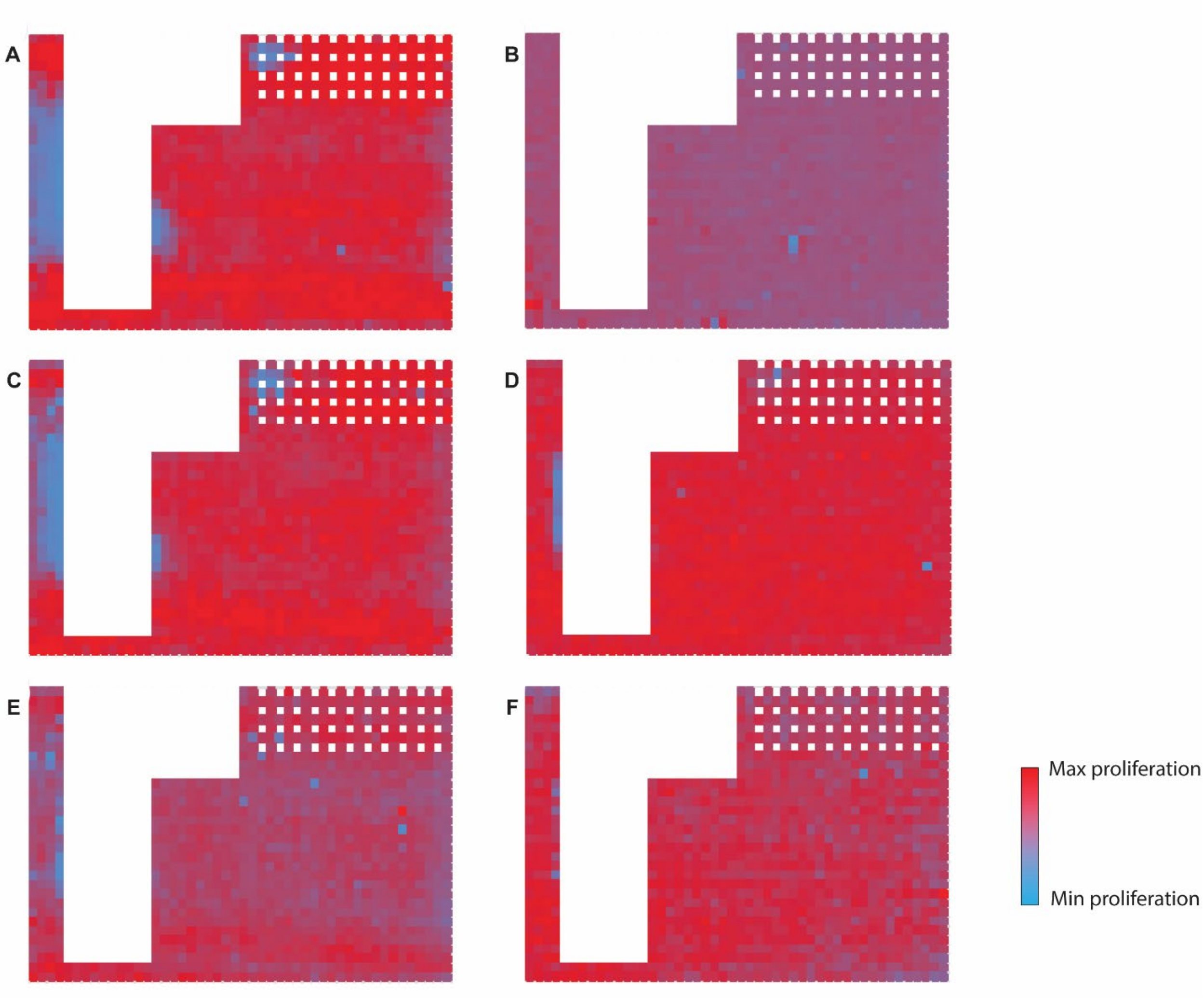


**Fig. S1. 1536-well assay optimization.** Shown are the optimization of 1536-well plate type and lid/seal to minimize compound leaching or effects of volatile compound on neighboring well. DMSO treated wells adjacent to compound treated wells (masked in white) were evaluated for modulation of parasite proliferation by SYBRGreen fluorescence after 72-h incubation; red is indicative of maximum parasite growth whereas blue indicates minimum growth. Color differential is maximized for each individual plate. **A)** polystyrene plate with light lid; **B)** cyclic olefin plate with foil seal; **C)** polystyrene plate heavy lid; **D)** cyclic olefin plate with heavy lid; **E)** polystyrene plate permeable seal; **F)** cyclic olefin plate with permeable seal. Illustrated are the leaching of artemether and artemisinin in the masked left hand wells to the DMSO control wells (not masked) and also in the single agent 1:3 dilution wells (right), diluted from left to right (top concentration 50nM), top compound artenimol (dihydroartemisinin), artemether, mefloquine and artemisinin. The polystyrene plates demonstrate a high degree of leaching (neighbor well effect) of compound that effects the growth of neighboring wells. There is a smaller degree of leaching in the cyclic olefin plates, which is further minimized through the use of breathable membrane seal **(F)**.

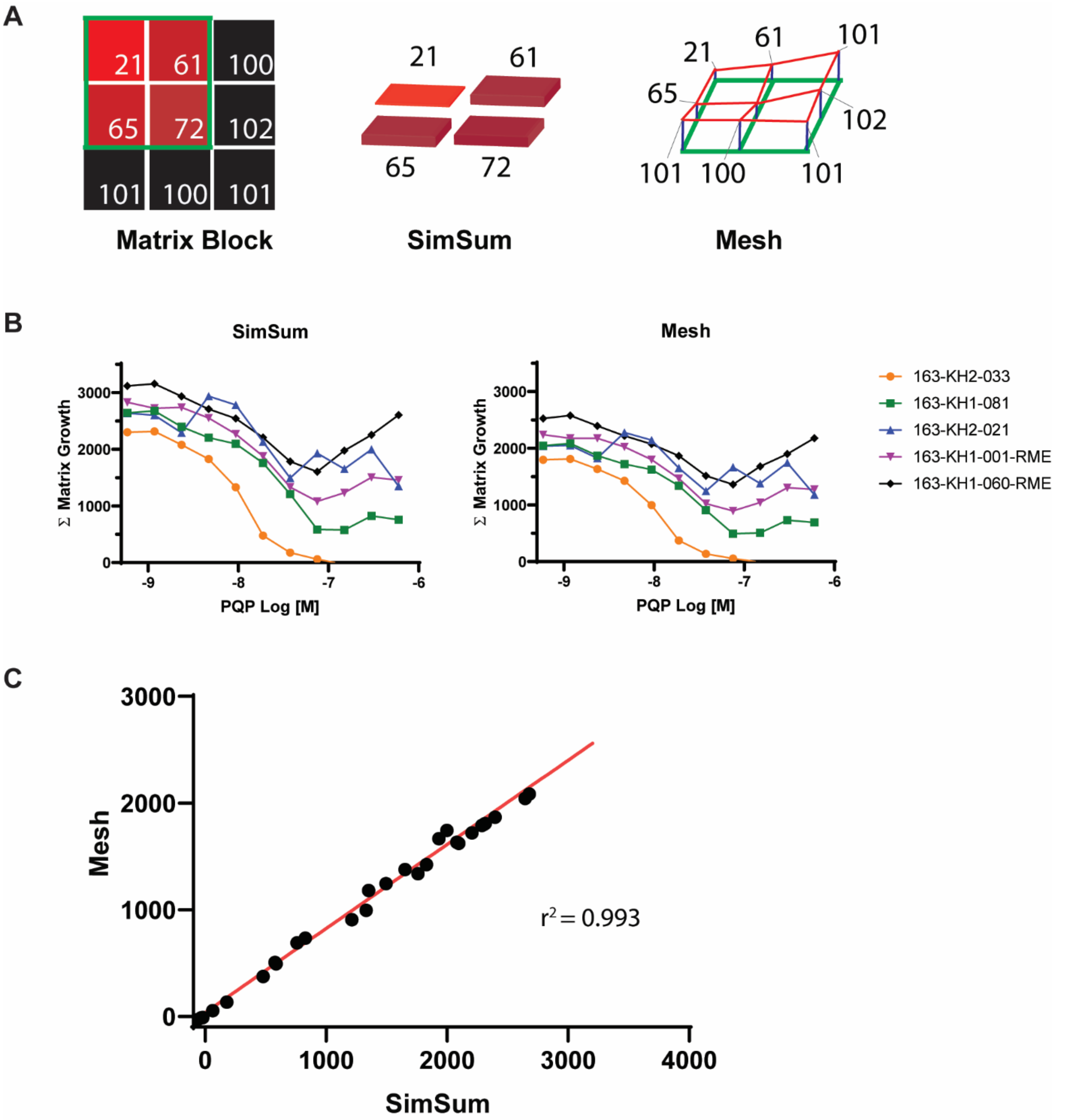


**Fig. S2. TACT calculation analysis methodology. A)** Representation of the Sum matrix analysis, represented in the 2-dimensional matrix, the simple sum (SimSum) where each block is constructed simply by extruding equally sized squares from their 2-dimensional log concentration grid locations and the Mesh method that is constructed by forming a vertex for each non-zero concentration pair in log-space and raising or lowering the vertex point in the third dimension relative to the measured response. **B)** Representative Sum matrix TACT dose response plots calculated both in the SimSum and Mesh methods. Overall, the pattern for each *P. falciparum* line is nearly equal between the two methods, although the Mesh method does have a decreased y-axis magnitude. **C)** Correlation plot between sum matrix calculated in both the SimSum and Mesh methods, $r^2$= 0.993 with a *P*-value <0.0001.

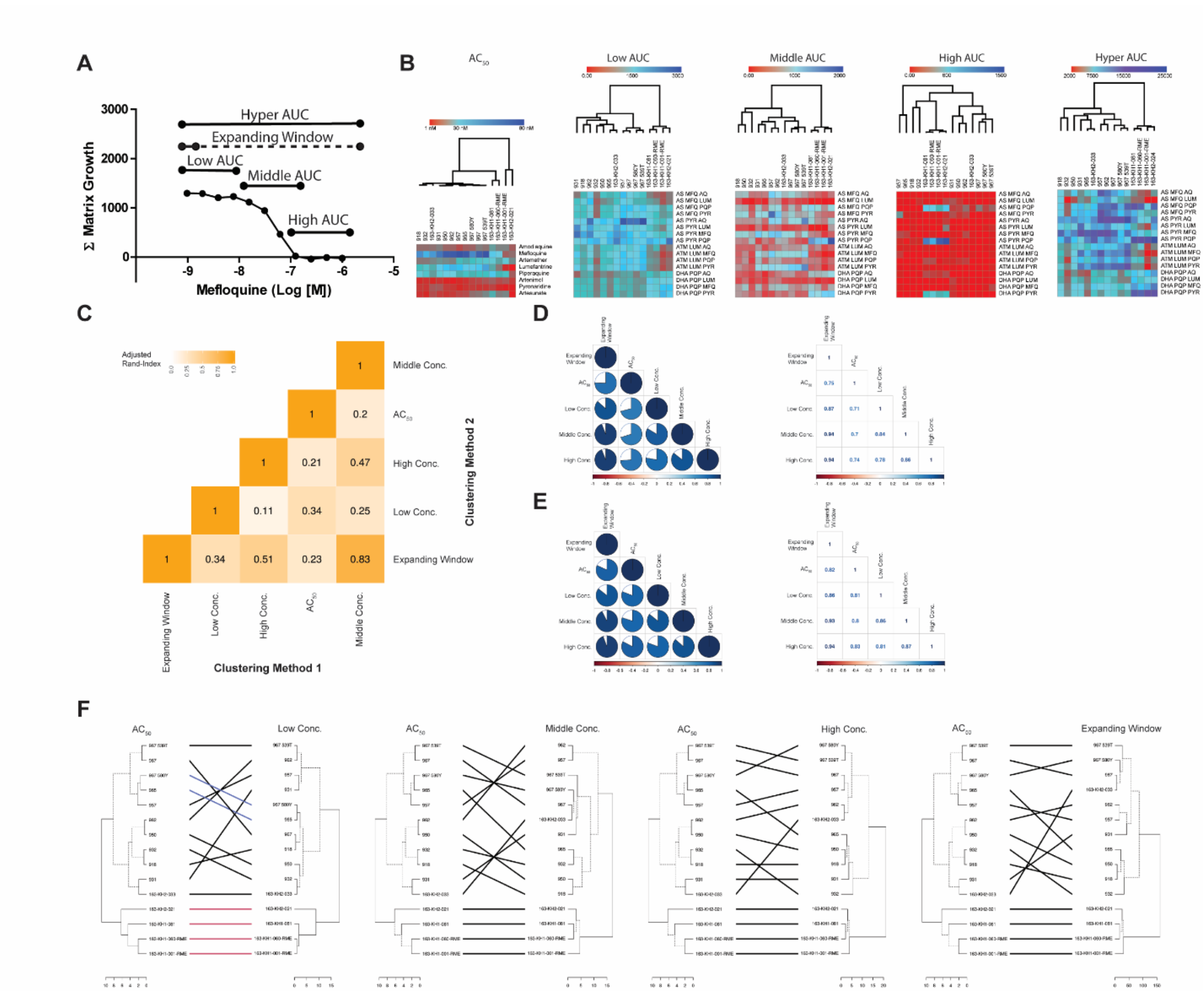


**Fig. S3. $AC_{50}$ - AUC Relationship Analysis.** Shown in panel **A)** are the subdivisions of the TACT response used for comparative analysis. The low area under the curve (AUC) division encompasses the area under the lowest 4 concentrations of drug C, middle AUC encompasses the next four concentration and the high AUC reflects the AUC under the highest concentration of drug C tested (overlaps with middle AUC on the fourth highest concentration point to have equal number of concentrations. The expanding-window is an increasing aggregation, from the lowest drug C concentration block, the sum of the lowest two drug C concentration blocks, the sum of the lowest three concentration blocks, *etc*. Hyper AUC reflects the AUC under the entire TACT response curve. (**B**) Qualitative comparison between the TACT responses for the lines tested, comparing the $AC_{50}$, low AUC, middle AUC, high AUC and hyper AUC, with hierarchical clustering based on response metrics (red reflects the minimum value and dark blue the maximum value). **C**) Pairwise adjusted Rand-index (ARI) of clustering produced by different clustering methods, white represents low agreement, whereas orange represents a high level of agreement. Quantitative correlation of dendrogram clustering, **D**) Cophenetic correlation, **E**) Baker correlation. **F**) Tanglegram dendrogram clustering of comparisons between $AC_{50}$ and low concentration AUC, middle concentration AUC, high concentration AUC and expanding-window.

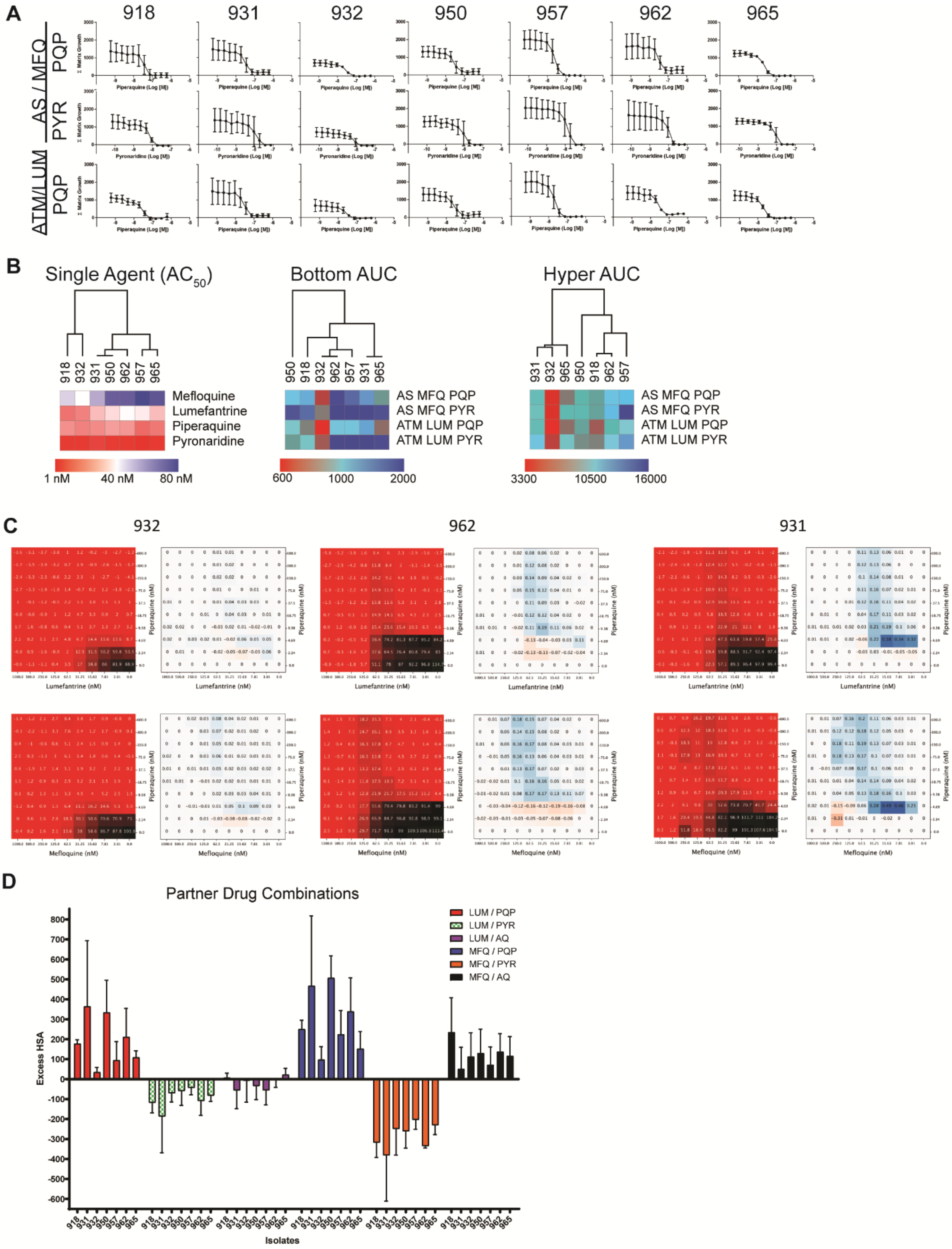


**Fig. S4. Drug interaction in 2009-2010 isolates. A**) TACT response curves for the 2009-2011 field isolates against artesunate/mefloquine/piperaquine, artesunate/mefloquine/pyronaridine and artemether/lumefantrine/piperaquine illustrating the differential activity against these lines, shown as mean ± standard deviation (n=3). **B**) Qualitative comparison between the TACT responses for the lines tested, comparing the single agent response ($AC_{50}$), low AUC and hyper AUC, with hierarchical clustering based on response metrics (red reflects the minimum value and dark blue the maximum value; tables S1 and S2). **C**) Representative normalized proliferation response for drug-drug (two compounds only) matrix assays, along with the excess HSA drug interaction plot, illustrating that different lines have variant drug-drug pharmacokinetic interactions. **D**) Excess HSA drug interaction model plot for lumefantrine/piperaquine (LUM/PQP), lumefantrine/pyronaridine (LUM/PYR), lumefantrine/amodiaquine (LUM/AQ), mefloquine/piperaquine (MFQ/PQP), mefloquine/pyronaridine (MFQ/PYR) and mefloquine/amodiaquine (MFQ/AQ) two-drug matrix assays, plotted as mean excess HSA ± standard deviation, n=3.

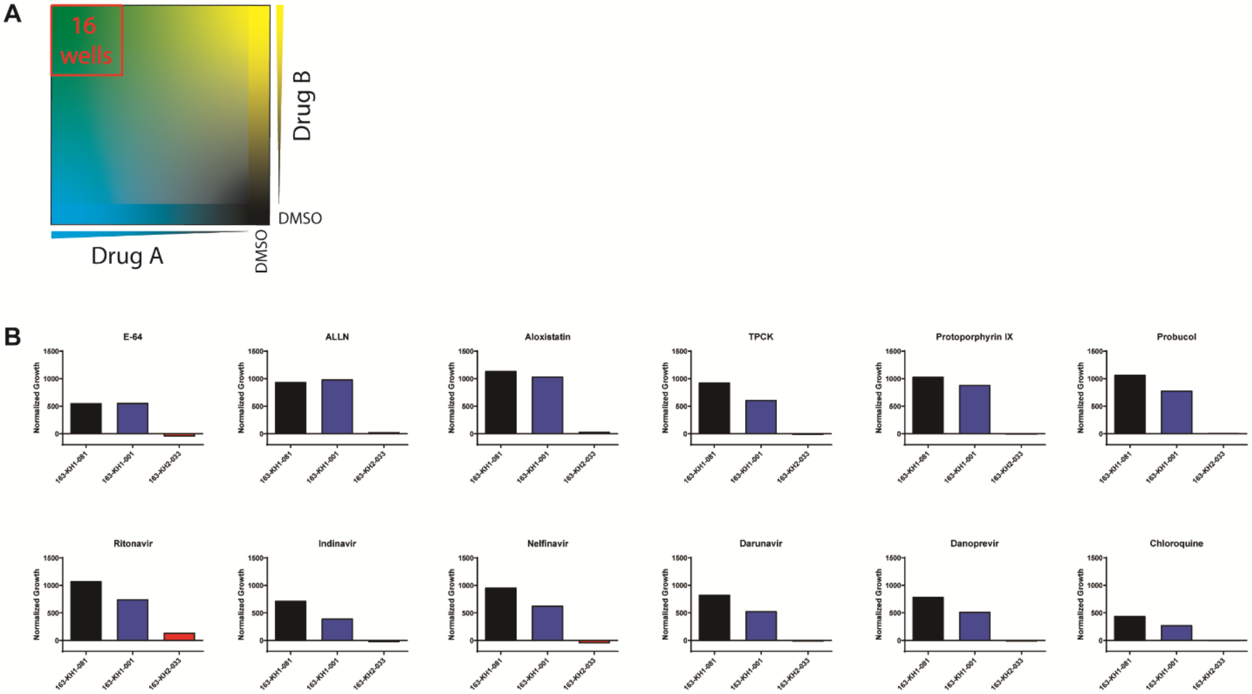


**Fig. S5. Screen for chemical inhibitors of PYR/PQP antagonism. A)** Since the antagonism between PYR and PQP is associated with parasites that have increased plasmepsin II/III copy number we questioned if this phenotype could be reversed by chemical means. Several protease inhibitors and other agents with potential activity within the parasite digestive vacuole were screened against PYR/PQP matrix blocks at four concentrations (either 10 or 1 µM top concentration, with serial 1:5 dilutions). **B)** Normalized growth for the 16 blocks with the top PYR/PQP concentrations were summed across all dilutions of the third agent. No inhibitor screened significantly modulated the PYR/PQP antagonism, however, slight antagonism was apparently induced in the 163-KH2-033 line with ritonavir.

### Table S1. Area under the triple artemisinin combination response curve

**High Area under the curve (High AUC) Triple combination response values**

| | 918 | 931 | 932 | 950 | 957 | 962 | 965 | 163-KH1-081 | 163-KH1-001-RME | 163-KH1-060-RME | 163-KH2-033 | 163-KH2-021 | 967 | 967 580Y | 967 539T |
|---|---|---|---|---|---|---|---|---|---|---|---|---|---|---|---|
| **AS MFQ AQ** | 57.8 | 33.0 | 22.7 | 13.9 | 39.2 | 27.9 | 39.6 | 50.8 | 61.7 | 51.0 | 29.2 | 23.6 | 63.7 | 57.9 | 33.0 |
| **AS MFQ LUM** | 40.2 | 195.8 | 120.1 | 132.2 | 62.4 | 50.5 | 52.7 | 74.2 | 50.9 | 33.3 | 70.4 | 164.6 | 41.5 | 29.0 | 82.9 |
| **AS MFQ PQP** | 17.4 | 145.6 | 7.8 | 159.5 | 8.8 | 261.2 | 11.7 | 694.8 | 422.5 | 658.2 | 112.3 | 725.0 | 289.2 | 157.1 | 235.2 |
| **AS MFQ PYR** | 71.3 | 182.7 | 55.5 | 121.1 | 349.9 | 164.0 | 90.0 | 33.4 | 49.8 | 75.9 | 16.2 | 112.0 | 24.5 | 75.0 | 53.2 |
| **AS PYR AQ** | 61.6 | 25.5 | 28.8 | 15.8 | 28.4 | 36.7 | 12.0 | 19.0 | 65.6 | 22.5 | 17.9 | 137.4 | 33.1 | 13.9 | 25.3 |
| **AS PYR LUM** | 80.1 | 53.3 | 46.6 | 29.9 | 24.0 | 8.1 | 39.6 | 48.8 | 22.1 | 12.6 | 28.0 | 305.1 | 20.4 | 75.1 | 108.2 |
| **AS PYR MFQ** | 54.1 | 75.3 | 32.0 | 59.6 | 179.9 | 174.3 | 147.6 | 43.8 | 13.1 | 85.7 | 62.3 | 31.8 | 37.7 | 33.0 | 79.3 |
| **AS PYR PQP** | 108.2 | 107.5 | 93.5 | 23.8 | 98.5 | 32.1 | 5.4 | 1021.0 | 1223.0 | 866.2 | 65.5 | 1405.0 | 21.1 | 47.2 | 66.0 |
| **ATM LUM AQ** | 61.7 | 75.3 | 47.0 | 61.9 | 74.8 | 65.5 | 88.3 | 90.4 | 73.6 | 22.1 | 46.6 | 16.5 | 7.2 | 70.5 | 79.4 |
| **ATM LUM MFQ** | 9.4 | 91.7 | 16.7 | 40.5 | 144.2 | 49.9 | 35.3 | 10.9 | 12.7 | 15.9 | 72.2 | 11.9 | 37.3 | 34.9 | 88.4 |
| **ATM LUM PQP** | 40.6 | 102.7 | 43.4 | 127.8 | 12.4 | 149.4 | 19.5 | 182.7 | 57.7 | 180.8 | 23.8 | 107.0 | 28.4 | 31.5 | 16.4 |
| **ATM LUM PYR** | 56.1 | 63.2 | 38.4 | 14.9 | 55.5 | 28.0 | 35.2 | 89.0 | 62.7 | 49.3 | 29.8 | 11.8 | 46.4 | 36.7 | 91.9 |
| **DHA PQP AQ** | 32.4 | 26.1 | 53.1 | 18.9 | 95.4 | 22.4 | 45.0 | 178.5 | 130.1 | 151.6 | 34.2 | 180.1 | 34.8 | 59.0 | 11.1 |
| **DHA PQP LUM** | 50.6 | 156.5 | 18.3 | 67.7 | 37.8 | 77.5 | 33.1 | 8.9 | 23.8 | 63.3 | 45.2 | 22.6 | 49.5 | 50.8 | 22.7 |
| **DHA PQP MFQ** | 17.0 | 115.3 | 17.3 | 44.9 | 72.3 | 51.1 | 37.7 | 25.9 | 42.3 | 30.9 | 30.3 | 48.3 | 42.2 | 81.2 | 173.2 |
| **DHA PQP PYR** | 17.0 | 118.7 | 46.0 | 65.0 | 22.4 | 84.9 | 48.1 | 449.0 | 415.0 | 539.1 | 37.6 | 454.5 | 64.8 | 41.1 | 8.8 |

**Middle Area under the curve (Middle AUC) Triple combination response values**

| | 918 | 931 | 932 | 950 | 957 | 962 | 965 | 163-KH1-081 | 163-KH1-001-RME | 163-KH1-060-RME | 163-KH2-033 | 163-KH2-021 | 967 | 967 580Y | 967 539T |
|---|---|---|---|---|---|---|---|---|---|---|---|---|---|---|---|
| **AS MFQ AQ** | 714.0 | 472.0 | 456.6 | 741.5 | 1173.0 | 1033.0 | 482.6 | 469.7 | 291.6 | 379.6 | 717.8 | 464.3 | 817.7 | 706.1 | 612.6 |
| **AS MFQ LUM** | 362.5 | 205.9 | 75.8 | 156.1 | 600.2 | 557.1 | 136.4 | 97.1 | 26.1 | 115.1 | 228.8 | 102.0 | 351.3 | 247.9 | 230.1 |
| **AS MFQ PQP** | 710.0 | 741.5 | 304.1 | 657.0 | 817.2 | 937.0 | 448.7 | 1096.0 | 554.2 | 849.1 | 425.5 | 883.7 | 714.7 | 456.8 | 906.0 |
| **AS MFQ PYR** | 808.8 | 966.2 | 399.6 | 888.4 | 1624.0 | 1223.0 | 908.5 | 822.3 | 424.1 | 530.9 | 662.8 | 405.1 | 797.2 | 634.1 | 682.2 |
| **AS PYR AQ** | 585.7 | 654.9 | 573.9 | 557.9 | 1122.0 | 1099.0 | 424.3 | 796.8 | 488.3 | 572.1 | 582.0 | 778.4 | 910.3 | 892.7 | 1480.0 |
| **AS PYR LUM** | 271.1 | 479.7 | 183.1 | 269.8 | 471.1 | 411.8 | 207.0 | 91.9 | 57.6 | 87.0 | 225.8 | 606.8 | 254.1 | 297.0 | 381.6 |
| **AS PYR MFQ** | 805.9 | 1187.0 | 684.6 | 824.1 | 1217.0 | 1247.0 | 869.6 | 530.4 | 220.2 | 472.5 | 904.0 | 369.6 | 863.4 | 561.8 | 876.4 |
| **AS PYR PQP** | 531.4 | 250.6 | 722.2 | 477.4 | 377.4 | 694.8 | 502.0 | 1411.0 | 1216.0 | 1249.0 | 659.9 | 1889.0 | 466.6 | 338.6 | 282.2 |
| **ATM LUM AQ** | 377.4 | 584.3 | 355.3 | 382.1 | 461.3 | 491.1 | 288.3 | 400.3 | 271.4 | 557.1 | 630.9 | 388.2 | 570.1 | 585.8 | 754.2 |
| **ATM LUM MFQ** | 369.4 | 961.4 | 273.5 | 542.6 | 1185.0 | 823.8 | 707.5 | 305.6 | 100.7 | 277.4 | 875.4 | 166.1 | 899.8 | 833.9 | 1126.0 |
| **ATM LUM PQP** | 410.9 | 720.2 | 232.1 | 584.5 | 645.4 | 643.3 | 375.9 | 485.9 | 281.6 | 475.2 | 459.2 | 423.8 | 473.1 | 375.3 | 553.2 |
| **ATM LUM PYR** | 572.8 | 1004.0 | 340.2 | 457.4 | 1083.0 | 1015.0 | 935.8 | 366.9 | 171.3 | 423.9 | 1075.0 | 350.8 | 1186.0 | 1205.0 | 1469.0 |
| **DHA PQP AQ** | 382.6 | 366.3 | 243.7 | 329.7 | 348.6 | 483.2 | 171.9 | 736.3 | 736.9 | 703.9 | 582.1 | 955.2 | 608.2 | 369.9 | 403.4 |
| **DHA PQP LUM** | 279.7 | 313.2 | 212.6 | 426.4 | 485.6 | 475.6 | 211.0 | 329.9 | 229.2 | 165.1 | 284.4 | 181.1 | 249.8 | 246.5 | 238.5 |
| **DHA PQP MFQ** | 593.8 | 706.0 | 328.9 | 758.7 | 712.9 | 758.2 | 372.2 | 765.5 | 371.7 | 664.0 | 756.6 | 665.2 | 582.0 | 502.3 | 622.2 |
| **DHA PQP PYR** | 593.8 | 515.9 | 375.6 | 528.6 | 285.1 | 490.2 | 242.9 | 1093.0 | 1184.0 | 963.9 | 408.6 | 1210.0 | 578.0 | 350.3 | 378.3 |

**Low Area under the curve (Low AUC) Triple combination response values**

| | 918 | 931 | 932 | 950 | 957 | 962 | 965 | 163-KH1-081 | 163-KH1-001-RME | 163-KH1-060-RME | 163-KH2-033 | 163-KH2-021 | 967 | 967 580Y | 967 539T |
|---|---|---|---|---|---|---|---|---|---|---|---|---|---|---|---|
| **AS MFQ AQ** | 1076.0 | 707.7 | 701.3 | 1197.0 | 1860.0 | 1601.0 | 1057.0 | 744.8 | 494.5 | 593.3 | 1192.0 | 722.3 | 1513.0 | 1327.0 | 1186.0 |
| **AS MFQ LUM** | 1409.0 | 1185.0 | 565.3 | 806.3 | 1632.0 | 1828.0 | 910.3 | 940.6 | 493.1 | 833.8 | 1163.0 | 715.7 | 1412.0 | 1049.0 | 1057.0 |
| **AS MFQ PQP** | 1154.0 | 1257.0 | 643.6 | 1174.0 | 1813.0 | 1468.0 | 1107.0 | 1485.0 | 909.5 | 1398.0 | 1485.0 | 1256.0 | 1802.0 | 1618.0 | 1757.0 |
| **AS MFQ PYR** | 1117.0 | 1186.0 | 601.3 | 1141.0 | 1825.0 | 1448.0 | 1149.0 | 1383.0 | 885.9 | 1130.0 | 1440.0 | 999.7 | 1639.0 | 1363.0 | 1642.0 |
| **AS PYR AQ** | 1630.0 | 1534.0 | 1642.0 | 1796.0 | 2629.0 | 2288.0 | 1649.0 | 1574.0 | 1109.0 | 1221.0 | 1787.0 | 1619.0 | 2292.0 | 2283.0 | 3141.0 |
| **AS PYR LUM** | 1600.0 | 2127.0 | 1357.0 | 1410.0 | 1749.0 | 1971.0 | 1529.0 | 1238.0 | 757.4 | 974.4 | 1982.0 | 1435.0 | 1592.0 | 1420.0 | 1572.0 |
| **AS PYR MFQ** | 1962.0 | 2297.0 | 1527.0 | 1622.0 | 1964.0 | 2144.0 | 1853.0 | 1565.0 | 1183.0 | 1299.0 | 2064.0 | 1362.0 | 1822.0 | 1506.0 | 1741.0 |
| **AS PYR PQP** | 1654.0 | 1478.0 | 1926.0 | 1507.0 | 1881.0 | 1792.0 | 1906.0 | 1935.0 | 1717.0 | 1757.0 | 2181.0 | 2279.0 | 1835.0 | 1662.0 | 1615.0 |
| **ATM LUM AQ** | 950.6 | 1111.0 | 786.2 | 1165.0 | 1615.0 | 1439.0 | 1173.0 | 843.7 | 545.2 | 1068.0 | 1346.0 | 737.5 | 1315.0 | 1580.0 | 1761.0 |
| **ATM LUM MFQ** | 1040.0 | 1697.0 | 802.5 | 1191.0 | 1797.0 | 1486.0 | 1390.0 | 831.8 | 389.6 | 743.5 | 1518.0 | 540.3 | 1563.0 | 1556.0 | 1838.0 |
| **ATM LUM PQP** | 941.0 | 1279.0 | 571.9 | 1153.0 | 1742.0 | 1212.0 | 1094.0 | 921.1 | 503.2 | 839.1 | 1624.0 | 657.6 | 1741.0 | 1631.0 | 1805.0 |
| **ATM LUM PYR** | 1061.0 | 1616.0 | 787.4 | 1202.0 | 1805.0 | 1536.0 | 1365.0 | 840.7 | 438.4 | 768.0 | 1576.0 | 642.0 | 1660.0 | 1623.0 | 1879.0 |
| **DHA PQP AQ** | 941.7 | 578.7 | 610.7 | 816.5 | 959.9 | 1037.0 | 688.9 | 1128.0 | 1247.0 | 960.2 | 871.1 | 1264.0 | 986.2 | 732.9 | 973.2 |
| **DHA PQP LUM** | 1040.0 | 1093.0 | 740.8 | 1086.0 | 960.8 | 1117.0 | 797.8 | 1667.0 | 1479.0 | 1346.0 | 1110.0 | 1548.0 | 1048.0 | 878.6 | 847.7 |
| **DHA PQP MFQ** | 1132.0 | 1167.0 | 737.4 | 1067.0 | 1043.0 | 1103.0 | 765.6 | 1540.0 | 1802.0 | 1388.0 | 1177.0 | 1810.0 | 1049.0 | 872.3 | 980.2 |
| **DHA PQP PYR** | 1132.0 | 1157.0 | 776.2 | 1076.0 | 891.3 | 1082.0 | 743.3 | 1551.0 | 1907.0 | 1423.0 | 1080.0 | 1706.0 | 1057.0 | 825.6 | 893.7 |

**Hyper Area under the curve (Hyper AUC) Triple combination response values**

| | 918 | 931 | 932 | 950 | 957 | 962 | 965 | 163-KH1-081 | 163-KH1-001-RME | 163-KH1-060-RME | 163-KH2-033 | 163-KH2-021 | 967 | 967 580Y | 967 539T |
|---|---|---|---|---|---|---|---|---|---|---|---|---|---|---|---|
| **AS MFQ AQ** | 7590.1 | 5059.4 | 4960.6 | 8511.9 | 13129.3 | 11375.6 | 6745.4 | 5101.3 | 3617.9 | 4075.9 | 8350.8 | 5274.8 | 10065.8 | 8832.6 | 7967.3 |
| **AS MFQ LUM** | 7691.2 | 5216.6 | 2293.8 | 3683.0 | 9611.3 | 10484.4 | 4471.5 | 4264.6 | 2083.6 | 4171.7 | 6407.3 | 2219.4 | 7640.8 | 5772.1 | 5267.7 |
| **AS MFQ PQP** | 8133.8 | 9223.8 | 4131.5 | 8579.1 | 11678.0 | 11376.9 | 6854.2 | 13686.6 | 7796.9 | 12112.6 | 8996.2 | 11881.2 | 12348.5 | 9703.8 | 12470.1 |
| **AS MFQ PYR** | 8114.1 | 9535.4 | 4127.7 | 8823.3 | 15517.4 | 11733.9 | 8894.9 | 9522.4 | 5580.2 | 7107.5 | 9395.2 | 5722.4 | 10658.1 | 9108.6 | 10571.5 |
| **AS PYR AQ** | 10216.3 | 9705.9 | 9905.5 | 10592.9 | 16569.5 | 14935.4 | 9380.4 | 10623.3 | 7264.0 | 7853.8 | 10739.8 | 11046.9 | 14250.2 | 14307.8 | 20545.7 |
| **AS PYR LUM** | 8126.0 | 11637.6 | 6769.7 | 7438.0 | 9787.5 | 10695.0 | 7662.6 | 5744.3 | 3583.5 | 4710.5 | 9967.2 | 9914.4 | 8170.8 | 7863.8 | 9031.1 |
| **AS PYR MFQ** | 12073.4 | 15618.9 | 9805.0 | 10854.2 | 14322.4 | 15355.8 | 12531.9 | 9415.3 | 6222.9 | 8062.8 | 13331.2 | 7592.7 | 11831.8 | 9103.7 | 11679.8 |
| **AS PYR PQP** | 9212.6 | 7404.5 | 11412.8 | 8949.5 | 9785.9 | 11253.0 | 10844.6 | 18284.5 | 17409.6 | 16308.5 | 12571.3 | 23204.0 | 10393.4 | 8917.7 | 8230.4 |
| **ATM LUM AQ** | 5640.5 | 7230.1 | 4953.8 | 6812.1 | 9259.3 | 8546.6 | 6238.0 | 5195.4 | 3349.1 | 7118.7 | 8664.1 | 4916.1 | 8503.7 | 10105.6 | 11597.9 |
| **ATM LUM MFQ** | 6208.2 | 11458.9 | 4781.8 | 7606.1 | 13190.1 | 10163.6 | 9348.4 | 5088.8 | 2111.0 | 4472.6 | 10695.3 | 3142.2 | 10806.5 | 10560.9 | 13067.0 |
| **ATM LUM PQP** | 5836.5 | 9156.4 | 3390.2 | 8109.9 | 10604.2 | 8694.8 | 6520.1 | 6833.8 | 3609.2 | 6443.8 | 9437.2 | 5132.0 | 10033.7 | 9072.6 | 10708.8 |
| **ATM LUM PYR** | 7100.7 | 11245.6 | 4929.2 | 7443.6 | 12359.2 | 10966.3 | 9856.6 | 5029.2 | 2523.1 | 5026.3 | 11315.8 | 4306.4 | 12053.3 | 12022.8 | 14302.8 |
| **DHA PQP AQ** | 5942.5 | 4092.6 | 3663.3 | 5136.2 | 5523.2 | 6642.2 | 3747.2 | 8721.5 | 9054.6 | 7712.6 | 6111.7 | 10167.2 | 6798.8 | 4629.3 | 6166.8 |
| **DHA PQP LUM** | 5676.0 | 5685.5 | 4234.9 | 6583.7 | 6315.0 | 6931.9 | 4461.9 | 9001.8 | 7558.3 | 6485.8 | 6143.1 | 7562.7 | 5643.2 | 4778.3 | 4744.6 |
| **DHA PQP MFQ** | 7495.2 | 7909.6 | 4711.6 | 7972.1 | 7800.3 | 7918.9 | 4913.1 | 10139.2 | 9731.5 | 8909.2 | 8434.6 | 11073.8 | 7103.8 | 6303.5 | 7703.8 |
| **DHA PQP PYR** | 7495.2 | 7204.3 | 5000.8 | 7312.1 | 5219.2 | 7224.0 | 4222.5 | 12870.9 | 14565.4 | 12073.8 | 6484.0 | 14016.5 | 6892.1 | 5060.9 | 5694.1 |

AS, artesunate; MFQ, mefloquine; AQ, amodiaquine; LUM, lumefantrine; PQP, piperaquine; PYR, pyronaridine; ATM, artemether; DHA, dihydroartemisinin

#### Table S2. Mean $AC_{50}$ values for single agent screening against *Plasmodium falciparum* lines tested

| | | Amodiaquine | Mefloquine | Artemether | Lumefantrine | Piperaquine | Artenimol | Pyronaridine | Artesunate |
|---|---|---|---|---|---|---|---|---|---|
| **918** | Mean $AC_{50}$* | 0.0112 | 0.0473 | 0.0122 | 0.0189 | 0.0237 | 0.0050 | 0.0063 | 0.0058 |
| | Standard deviation | 0.0022 | 0.0287 | 0.0066 | 0.0097 | 0.0032 | 0.0031 | 0.0043 | 0.0028 |
| | Independent replicates | 7 | 7 | 7 | 7 | 7 | 7 | 7 | 7 |
| **931** | Mean $AC_{50}$ | 0.0128 | 0.0530 | 0.0149 | 0.0309 | 0.0232 | 0.0059 | 0.0094 | 0.0066 |
| | Standard deviation | 0.0066 | 0.0313 | 0.0097 | 0.0172 | 0.0072 | 0.0051 | 0.0066 | 0.0023 |
| | Independent replicates | 6 | 6 | 6 | 5 | 6 | 6 | 6 | 6 |
| **932** | Mean $AC_{50}$ | 0.0112 | 0.0402 | 0.0122 | 0.0214 | 0.0214 | 0.0056 | 0.0059 | 0.0048 |
| | Standard deviation | 0.0020 | 0.0159 | 0.0063 | 0.0122 | 0.0061 | 0.0044 | 0.0038 | 0.0021 |
| | Independent replicates | 7 | 7 | 7 | 7 | 7 | 7 | 7 | 7 |
| **950** | Mean $AC_{50}$ | 0.0104 | 0.0654 | 0.0139 | 0.0353 | 0.0262 | 0.0044 | 0.0066 | 0.0058 |
| | Standard deviation | 0.0030 | 0.0266 | 0.0038 | 0.0259 | 0.0092 | 0.0024 | 0.0061 | 0.0023 |
| | Independent replicates | 7 | 7 | 7 | 7 | 7 | 7 | 7 | 7 |
| **957** | Mean $AC_{50}$ | 0.0081 | 0.0797 | 0.0185 | 0.0374 | 0.0174 | 0.0050 | 0.0048 | 0.0076 |
| | Standard deviation | 0.0017 | 0.0359 | 0.0075 | 0.0192 | 0.0035 | 0.0012 | 0.0037 | 0.0015 |
| | Independent replicates | 7 | 7 | 7 | 7 | 7 | 7 | 7 | 7 |
| **962** | Mean $AC_{50}$ | 0.0104 | 0.0662 | 0.0132 | 0.0399 | 0.0252 | 0.0034 | 0.0062 | 0.0063 |
| | Standard deviation | 0.0034 | 0.0267 | 0.0055 | 0.0251 | 0.0067 | 0.0017 | 0.0050 | 0.0021 |
| | Independent replicates | 7 | 7 | 7 | 7 | 7 | 7 | 7 | 7 |
| **965** | Mean $AC_{50}$ | 0.0100 | 0.0693 | 0.0171 | 0.0308 | 0.0200 | 0.0057 | 0.0058 | 0.0081 |
| | Standard deviation | 0.0035 | 0.0656 | 0.0052 | 0.0157 | 0.0045 | 0.0023 | 0.0035 | 0.0032 |
| | Independent replicates | 7 | 7 | 7 | 7 | 7 | 7 | 7 | 7 |
| **163-KH1-001-RME** | Mean $AC_{50}$ | 0.0133 | 0.0098 | 0.0234 | 0.0054 | 0.0203 | 0.0091 | 0.0037 | 0.0137 |
| | Standard deviation | 0.0028 | 0.0049 | 0.0159 | 0.0040 | 0.0131 | 0.0050 | 0.0018 | 0.0190 |
| | Independent replicates | 7 | 7 | 7 | 7 | 7 | 7 | 6 | 7 |
| **163-KH1-060-RME** | Mean $AC_{50}$ | 0.0132 | 0.0270 | 0.0165 | 0.0120 | 0.0182 | 0.0080 | 0.0051 | 0.0129 |
| | Standard deviation | 0.0029 | 0.0127 | 0.0063 | 0.0084 | 0.0079 | 0.0064 | 0.0026 | 0.0103 |
| | Independent replicates | 6 | 6 | 6 | 6 | 6 | 6 | 6 | 6 |
| **163-KH1-081** | Mean $AC_{50}$ | 0.0156 | 0.0312 | 0.0231 | 0.0169 | 0.0331 | 0.0101 | 0.0072 | 0.0176 |
| | Standard deviation | 0.0046 | 0.0145 | 0.0204 | 0.0174 | 0.0220 | 0.0076 | 0.0058 | 0.0261 |
| | Independent replicates | 6 | 6 | 6 | 6 | 6 | 6 | 6 | 6 |
| **163-KH2-021** | Mean $AC_{50}$ | 0.0097 | 0.0114 | 0.0142 | 0.0029 | 0.0095 | 0.0066 | 0.0037 | 0.0035 |
| | Standard deviation | 0.0036 | 0.0008 | 0.0052 | 0.0005 | 0.0077 | 0.0040 | 0.0020 | 0.0022 |
| | Independent replicates | 3 | 3 | 3 | 3 | 3 | 3 | 3 | 3 |
| **163-KH2-033** | Mean $AC_{50}$ | 0.0116 | 0.0451 | 0.0136 | 0.0173 | 0.0157 | 0.0046 | 0.0092 | 0.0082 |
| | Standard deviation | 0.0015 | 0.0215 | 0.0045 | 0.0088 | 0.0018 | 0.0003 | 0.0057 | 0.0023 |
| | Independent replicates | 3 | 3 | 3 | 3 | 3 | 3 | 3 | 3 |
| **967** | Mean $AC_{50}$ | 0.0128 | 0.0617 | 0.0211 | 0.0338 | 0.0191 | 0.0077 | 0.0058 | 0.0120 |
| | Standard deviation | 0.0049 | 0.0279 | 0.0071 | 0.0214 | 0.0035 | 0.0030 | 0.0044 | 0.0074 |
| | Independent replicates | 7 | 7 | 7 | 7 | 7 | 7 | 7 | 7 |
| **967 580Y** | Mean $AC_{50}$ | 0.0102 | 0.0484 | 0.0191 | 0.0313 | 0.0179 | 0.0071 | 0.0066 | 0.0113 |
| | Standard deviation | 0.0030 | 0.0240 | 0.0092 | 0.0199 | 0.0061 | 0.0058 | 0.0030 | 0.0069 |
| | Independent replicates | 7 | 7 | 7 | 7 | 7 | 7 | 7 | 7 |
| **967 539T** | Mean $AC_{50}$ | 0.0117 | 0.0735 | 0.0219 | 0.0434 | 0.0192 | 0.0089 | 0.0061 | 0.0137 |
| | Standard deviation | 0.0028 | 0.0439 | 0.0095 | 0.0310 | 0.0046 | 0.0041 | 0.0035 | 0.0080 |
| | Independent replicates | 7 | 7 | 7 | 7 | 7 | 7 | 7 | 7 |

Mean $AC_{50}$ values are expressed in µM, derived from 72 hr SYBRGreen-1 in vitro proliferation assays

#### Table S3. Entanglement of dendrograms generated by different clustering methods

| Clustering Method 1 | Clustering Method 2 | Entaglement |
|---|---|---|
| Single Agent ($AC_{50}$) | TACT AUC- Low Conc. | 0.22 |
| Single Agent ($AC_{50}$) | TACT AUC- Middle Conc. | 0.14 |
| Single Agent ($AC_{50}$) | TACT AUC- High Conc. | 0.13 |
| Single Agent ($AC_{50}$) | TACT AUC- Sliding Window | 0.26 |

$AC_{50}$, single agent drug activity that inhibits parasite proliferation by 50% *in vitro* ; TACT, triple artemisinin conbination therapy; AUC, area under the curve; Conc., concentration

#### Table S4. Parasite genotypes

| | 918 | 931 | 932 | 950 | 957 | 962 | 965 | 967 | 163-KH1-081 | 163-KH1-001-RME | 163-KH1-060-RME | 163-KH2-033 | 163-KH2-021 |
|---|---|---|---|---|---|---|---|---|---|---|---|---|---|
| PfMDR1 | N86 | N86 | N86 | N86 | N86 | N86 | N86 | N86 | N86 | N86 | N86 | N86 | N86 |
| | **Y184F** | **Y184F** | Y184 | Y184 | **Y184F** | Y184 | **Y184F** | **Y184F** | **Y184F** | **Y184F** | **Y184F** | Y184 | **Y184F** |
| | D1246 | D1246 | D1246 | D1246 | D1246 | D1246 | D1246 | D1246 | D1246 | D1246 | D1246 | D1246 | D1246 |
| Pfmdr1 CNV | Single | Single | Single | **3 copies** | **3 copies** | **3 copies** | **2 copies** | **2 copies** | Single | Single | Single | Single | Single |
| PfCRT | **K76T** | **K76T** | **K76T** | **K76T** | **K76T** | **K76T** | **K76T** | **K76T** | **K76T** | **K76T** | **K76T** | **K76T** | **K76T** |
| | H97 | H97 | H97 | H97 | H97 | H97 | H97 | H97 | H97 | H97 | **H97Y** | H97 | **H97Y** |
| | **A220S** | **A220S** | **A220S** | **A220S** | **A220S** | **A220S** | **A220S** | **A220S** | **A220S** | **A220S** | **A220S** | **A220S** | **A220S** |
| | **Q271E** | **Q271E** | **Q271E** | **Q271E** | **Q271E** | **Q271E** | **Q271E** | **Q271E** | **Q271E** | **Q271E** | **Q271E** | **Q271E** | **Q271E** |
| | **N326S** | **N326S** | **N326S** | **N326S** | **N326S** | **N326S** | **N326S** | **N326S** | **N326S** | **N326S** | **N326S** | **N326S** | **N326S** |
| | M343 | M343 | M343 | M343 | M343 | M343 | M343 | M343 | M343 | **M343L** | M343 | M343 | M343 |
| | **I356T** | **I356T** | **I356T** | **I356T** | **I356T** | **I356T** | **I356T** | **I356T** | **I356T** | **I356T** | **I356T** | **I356T** | **I356T** |
| | A366 | A366 | A366 | A366 | A366 | A366 | A366 | A366 | **A366T** | A366 | A366 | A366 | A366 |
| | **R371I** | **R371I** | **R371I** | **R371I** | **R371I** | **R371I** | **R371I** | **R371I** | **R371I** | **R371I** | **R371I** | **R371I** | **R371I** |
| Kelch13 | R539 | R539 | R539 | R539 | **R539T** | R539 | R539 | **R539T** | R539 | R539 | R539 | R539 | R539 |
| | Y493 | Y493 | Y493 | **Y493H** | Y493 | **Y493H** | **Y493H** | Y493 | Y493 | Y493 | Y493 | Y493 | Y493 |
| | C580 | C580 | C580 | C580 | C580 | C580 | C580 | C580 | **C580Y** | **C580Y** | **C580Y** | C580 | **C580Y** |
| Plasmepsin II/III CNV | Single | Single | Single | Single | Single | Single | Single | Single | **Amplified** | **Amplified** | **Amplified** | Single | **Amplified** |

PfMDR1, *Plasmodium falciparum* multdrug resistance transporter 1 (PF3D7_0523000); PfCRT, *P. falciparum* chloroquine resistance transporter (PF3D7_0709000), all lines have the CVIET allele for PfCRT positions 72-76; Kelch13 (PF3D7_1343700); Plasmepsin II/III copy number variation (PF3D7_1408000/PF3D7_1408100). (*11,57*)

#### Table S5. *Plasmodium falciparum* SYBRGreen assay 1,536-well protocol

| Step | Parameter | Value | Description |
|---|---|---|---|
| 1 | Reagent | 3 µL | Complete Malaria Growth medium |
| 2a<br>2b | Library Compounds<br>Control Compounds | 20 nL<br>20 nL | Compound source plates<br>Artesunate 120 µM (final assay concentration 300 nM) |
| 3 | Reagent | 5 µL | Malaria-infected RBCs |
| 4 | Seal | | Breath-easy Sealing Membrane |
| 5 | Time | 72 hr | 37°C incubation (5% $O_2$, 5% $CO_2$, 90% $N_2$; 95% $R_h$) |
| 6 | Peel | | Seals removed with X-Peel automated microplate seal remover |
| 7 | Reagent | 2 µL | Lysis buffer + SYBRGreen |
| 8 | Time | Overnight | 37°C incubation (5% $O_2$, 5% $CO_2$, 90% $N_2$; 95% $R_h$) |
| 9 | Detector | Fluorescence | EnVision |

| Step | Notes |
|---|---|
| 1 | Assay adopted from Plouffe *et al*. 2008 PNAS 105: 9059. Reagents were dispensed into 1536-well black clear Cyclo-Olefin Polymer plate (Aurora Microplates) using a Multidrop Combi (Thermo Fisher Scientific Inc.) contained in a biosafety cabinet. |
| 1a | Complete Malaria Growth Media: 1X RPMI-1640 w/L-glutamine, 5.5g/L Albumax II, 367 µM Hypoxanthine, 25 mM HEPES, 0.25% Sodium Bicarbonate, 35 µM Gentamicin sulfate |
| 2 | Compound source plates. Single acoustic ejection dispense. |
| 3 | Hematocrit 4% (final 2.5%), 0.3% parasitemia |
| 5 | Lysis buffer: 20 mM Tris-HCl, 10 mM EDTA, 0.16% Saponin, 1.6% Triton-X, 10X SYBR Green.<br>25 sec shake immediately following dispense |
| 8 | Approximately 18 hours |
| 9 | EnVision (PerkinElmer) bottom read at 485/14nm excitation and 535/25 nm emission |
